# ART: A Compositional Physical Theory for the End-to-End Design of Solid-Phase Immunosensors

**DOI:** 10.64898/2026.10.27.762221

**Authors:** Jinghui Zhang, Tianyuan Zhao, Hao Li, Xiaotian Tan

## Abstract

Predicting complete solid-phase immunoassay performance remains challenging because antibody recognition, reactor transport, molecular transduction and physical observation are coupled rather than independent. Here, we formulate ART as a compositional physical theory expressed as Y = T_s_[T_m_[AR(X)]]. Recognition A and reactor transport R jointly determine the retained molecular state, while molecular transduction T_m_ and physical observation T_s_ operate through replaceable interfaces. Finite-time kinetic and reaction–diffusion calculations across plate, tubular and bead formats show that antibodies with identical dissociation constants can produce different capture responses because of distinct association and dissociation rates. In fixed-length tubular reactors, mass-conserving transport distinguishes surface capture density from fractional recovery, revising a first-passage-based local-density optimum while preserving a multicriterion design window. An AR-only concentration-threshold map identifies when recognition kinetics or reactor redesign offers greater local leverage. The benefits of improved affinity also depend on the kinetic pathway and reactor regime. For enzymatic transduction, we propose a four-parameter, substrate-conserving model in which the saturation-shape parameter is h_m_ = K_M_/S_0_ under a one-effective-substrate approximation. Accumulated TMB color and time-dependent luminol emission represent distinct consequences of reaction progress, while Ts describes measurement limits through a simplified floor–slope–roof model. ELISA with OD450 readout and tubular chemiluminescence with camera detection illustrate the end-to-end framework. These examples remain conditional model demonstrations, and existing complete-chain curves have not yet been recalibrated using the new enzyme model. ART provides a modular quantitative framework for assay design based on explicit physical interfaces and coupled processes.

## Introduction

### A mature experimental field, but a fragmented quantitative language

For decades, solid-phase immunoassays have served as central tools in molecular diagnostics and biological research. Conventional microwell ELISA, microfluidic surface assays, bead-based immunoassays and single-molecule digital formats have evolved along different engineering routes, each improving an important dimension of analytical performance. Tubular tip optofluidic immunoassay (TOI) is a concrete implementation of this evolution: the published hLife study demonstrated binding- and neutralizing-antibody analysis using a microliter-scale fingertip specimen.[1] The earlier first-passage-time (FPT) study then attempted to quantify the role of molecular encounter kinetics and tube size; its physical design questions provide an important starting point for the more complete AR analysis below.[2] Better antibodies improve molecular recognition; smaller reaction spaces and mixing schemes change transport and incubation time; enzyme and fluorophore labels increase the physical reporter output; and increasingly capable optical or electronic instruments resolve weaker signals. The resulting experimental landscape is rich, but a practical theoretical difficulty persists: how should one predict the performance of a complete immunoassay from the independently measured properties of its components?

This difficulty becomes clear when comparing what researchers usually measure. Antibodies are often ranked by K_D_ or paired k_on_/k_off_; reactors by dimensions, sample volume or capture efficiency; reporter chemistries by catalytic activity, quantum yield or accumulated signal; and detectors by responsivity, quantum efficiency, exposure time, noise floor or dynamic range. Yet the final assay is judged by concentration response, analytical specificity, limit of detection (LOD), quantification range, sample requirement and assay time. These quantities do not share a common denominator. An antibody with lower K_D_ need not deliver more molecules to a surface in a short assay; a reactor that captures a larger fraction of a small sample need not maximize bound molecules per unit area; greater chemical amplification does not distinguish target complexes from cross-reactive ones; and lower camera read noise cannot recreate photons that the optical system never collects. The challenge is to explain and predict these differences within one assay, not to rank components in isolation.

There is substantial prior theory on which such a description should build. Molecular mass-action kinetics, surface reaction–diffusion and transport-limited capture have long been modeled, including for microfluidic assays and surface-binding sensors. Previous analyses have also treated the quantitative consequences of nonspecific binding, finite molecular counting statistics and digital immunoassay architectures. These contributions are essential rather than superseded by ART. They establish that transport, binding, background and detection are each consequential, but they often address distinct platforms, limiting regimes or interfaces. A remaining practical need is an operational, cross-platform way to connect independently characterized component models to the final measured signal and its observability constraints, without assuming ideal chemistry or conflating chemical sensitivity with instrumental sensitivity.[6–12]

### Why optimizing isolated components does not determine assay performance

Consider four seemingly straightforward design questions. Does a tenfold lower K_D_ imply a tenfold improvement in assay sensitivity? Does a tenfold reduction in reactor dimension necessarily produce a stronger local signal? Does adding PolyHRP improve LOD in proportion to enzymatic amplification? Does a lower instrumental detection floor ensure that an observed event is chemically specific? In general, none has a platform-independent answer. Their answers depend on the finite-time sequence through which molecules are sampled, transported, captured, retained, labeled, converted into a physical reporter and finally observed in the presence of target-derived and background fluctuations.

Three familiar analytical simplifications expose the problem. First, a device-oriented analysis may effectively assume perfect recognition, allocating all prospective improvement to geometry, amplification or detection. Second, a confinement-oriented analysis may equate increased molecular arrival or recovered fraction with increased analytical performance, without first specifying bound surface density or downstream observation. Third, an ultra-sensitive detection analysis may treat the ability to register every reporter event as equivalent to identifying a true molecular target. Each approximation is useful in its own limit, but none is sufficient as a general endpoint-assay model. These three idealizations are used here as explicit thought experiments, not as established labels or descriptions of whole research communities. Their limits are revisited in the Discussion.

A second obstacle is the standard use of fitted calibration curves as substitutes for causal models. A logistic or other empirical response curve can summarize an assay after it has been built and calibrated, but it cannot by itself explain why changing k_off_, tube radius, washing time, HRP loading or exposure duration changes the curve. Even a mechanistic model remains incomplete if it predicts only retained antibody complexes but is then interpreted as a prediction of OD450, collected photons or analytical LOD. What is needed is a mathematical language that passes well-defined physical quantities, with explicit units and operating limits, from one stage into the next.

### From attempted decoupling to the AR–T construction

We therefore begin by asking which functions can legitimately be separated. A denotes independently measurable recognition inputs: binding and unbinding kinetics, cross-reactivity, non-specific interactions, surface accessibility and buffer dependence. R denotes the experimental boundary-value problem: geometry, diffusivity, liquid inventory, available surface sites, mixing, incubation, washing and the specified endpoint. These are separable parameter sets and can be perturbed in controlled experiments. They are not, however, two independent sequential operations that each yield an observable immobilized molecular population. The wall concentration depends on transport and binding history; the reaction flux simultaneously depletes the liquid and changes surface occupancy. For a solid-phase endpoint assay, the first physical state-generating map is therefore the coupled AR process, whose output is a population or distribution of retained target, off-target and non-specifically reporter-competent complexes.

The output of AR also identifies the useful boundary between chemical recognition and transduction. From a known reporter-competent population, molecular transduction T_m_ describes how reaction or labeling chemistry creates the next physical signal. HRP–TMB develops an accumulating chromophore; HRP–luminol produces photons in real time with finite chemical yield; fluorescent labels cycle through excitation and photon emission with efficiency and photobleaching constraints. Amplification, including PolyHRP loading or high-yield labels, belongs here. By contrast, physical observation T_s_ describes the later conversion from that physical signal to what is collected and recorded: optical path, absorption wavelength, collection solid angle, exposure, quantum efficiency, detector noise, saturation and reporting rules. T_m_ and T_s_ are thus modeled as separate conditional mappings because their interface can be specified by a physical reporter state; separation does not imply that optimization of one is independent of constraints imposed by the other.

These distinctions lead to the defining compositional architecture

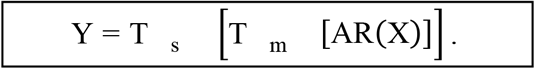

The notation defines physical interface order and does not prescribe a single formula for every transduction chemistry. Here X describes the specimen, AR supplies retained true and false reporter-competent populations, T_m_ produces a chemically defined reporter state, and T_s_ maps that state to the reported observation. The primary working models below use effective deterministic transfer functions and a conservatively chosen detection floor; specialized stochastic extensions are optional rather than required for the core composition. The scientific contribution lies in identifying the admissible coupling and modularity rules, not in claiming that composing functions or using diffusion equations is new mathematics. A and R are independently measurable design inputs but not two independent surface-population transfer functions; the AR output must be generated by a joint mass-conserving boundary problem. The AR-to-T_m_ and T_m_-to-T_s_ interfaces, in contrast, are defined by distinguishable molecular and physical signal states. Once their units, relevant time windows and effective operating conditions are specified, different models may be substituted within a module and subjected to prospective predictions without re-defining ART itself. These physical interfaces and their coupling relationships are summarized in Figure 1.

**Figure 1.**
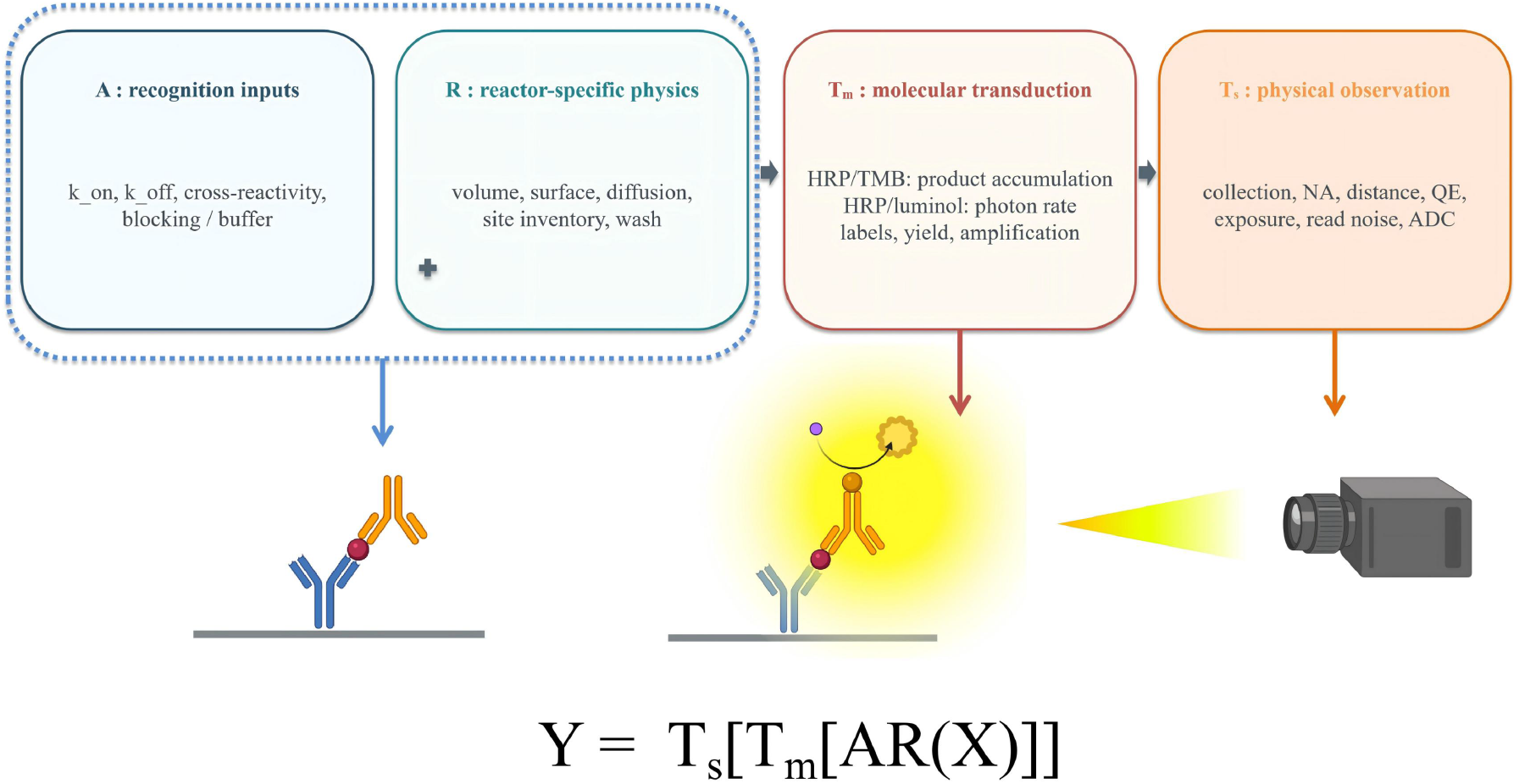
ART requires physically justified interfaces, not a universal kinetic law. Recognition kinetics A and reactor R can be measured separately, yet the bound populations arise from one coupled AR model. Molecular conversion T_m_ and reported observation T_s_ follow as replaceable, physically typed interfaces.

### A theory of coupling rules rather than a privileged set of component equations

The architecture makes a claim about how physical subproblems communicate. Recognition chemistry alone specifies microscopic rates and competing binding channels, but not a unique finite-time surface population. Transport and reversible adsorption are coupled through the same wall flux; they must be solved as AR, regardless of whether the implementation uses a spatial PDE, a validated reduced model or another conservative numerical method. Thus replacing the AR solver is permissible, while replacing the physically coupled AR process with a free-standing affinity-to-signal gain is not.

By contrast, the retained molecular state can serve as a common input to alternative molecular reporter modules, such as HRP–TMB product formation, fluorescence under illumination or time-dependent luminol emission. Their outputs are chemically and temporally distinct but may be passed onward to a corresponding observation module that accounts for wavelength, sample geometry, collection, exposure, quantum efficiency and electronic noise. A valid module replacement preserves the meaning, units, timing and conservation or saturation limits of the exchanged physical state. Molecular recognition background remains present in downstream outputs; a reported signal beneath the observation floor is set to zero under the operational rule used here. This concise working description does not require a separate detector-noise calculation.

This leads to two complementary standards of success. First, within-module predictions must withstand equation-level checks (mass conservation, limiting cases, numerical convergence and independently determined parameters). Second, end-to-end predictions must be tested without silently refitting the interfaces to each new assay. The first standard is most developed for AR in this manuscript; the T instantiations demonstrate the compositional interfaces while retaining chemistry-specific open calibration questions. Consequently the theoretical framework is broader than the present collection of numerical examples, but its experimentally supported scope must not be overstated.

### What the theory must predict, and how it is tested here

A useful theory should do more than classify assay components. It should (i) predict the concentration-to-signal transfer function under independently specified chemistry, geometry and readout; (ii) identify which intervention has the greatest leverage at a given operating point; and (iii) support inverse design under constraints on LOD, specificity, sample volume, time and upper measurement range. These three tasks—forward prediction, limitation attribution and constrained design—provide the organizing questions for the rest of the paper.

We first treat A at a fixed reference reactor, using same-K_D_ kinetic contrasts and human/goat IgG recognition to distinguish intrinsic affinity from finite-time selectivity. Uniform-rate perturbation hypotheses for casein and PBST show how apparently indiscriminate chemistry can nevertheless alter true and false binding unequally. We next derive the coupled AR surface/transport problem, retain its mathematical approximations and applicability limits, and compare kinetic families across well, tubular and bead geometries without confusing surface density, total captured molecules and recovered fraction. For the tubular reactor we examine finite molecular inventory, affinity-improvement pathways, tube radius, and time-to-80%-equilibrium (t_80_) as distinct design observables; the moment-informed FPT exponential approximation is explicitly checked against the relevant diffusion boundary problem. We then separate T_m_ from T_s_ using a substrate-conserving four-parameter conversion law, accumulation versus instantaneous-emission outputs, and a minimal effective floor–slope–roof observation operator. Finally, independent ELISA– HRP/TMB/OD450 and TOI–HRP/luminol/camera examples complete the forward chain with a shared illustrative A but distinctly modeled reactors.

The present scope is deliberately specific: finite-time, washed or endpoint solid-phase immunoassays, especially optical readouts. It does not claim universal coverage of continuous real-time sensors, electrochemical transducers or every binding stoichiometry. Reported experimental observations, published physical constants, exact limits, reduced-model calculations and hypothetical instrument parameters are distinguished throughout. The central proposition to be tested is not whether each individual physical principle is known, but whether its compositional use yields transferable, prospective predictions that cannot be obtained from component-only intuition.

### Recognition (A): kinetics and selectivity under a defined reactor

Question: Can molecular recognition be ranked by one affinity constant when binding duration, reagent supply and background channels matter? The following controlled A calculations establish the kinetic variables that AR must subsequently carry into a physical reactor.

### Why KD alone is insufficient

The intrinsic equilibrium affinity K_D_ = k_off_/k_on_ specifies the balance of association and dissociation; it does not specify how quickly finite-time binding approaches that equilibrium. This distinction is particularly important under short incubation, finite reagent supply and a subsequent wash. A defensible “A comparison” immobilizes the comparison protocol: for a reference reactor R_0_, vary a while holding geometry, accessible capacity, incubation and wash fixed. The observed binding is still AR a, R_0_, not an autonomous A X product.

In a well-mixed, reversible finite-bath reference with binding capacity B_0_ and initial soluble ligand C_0_, a simple surface-binding step satisfies dx

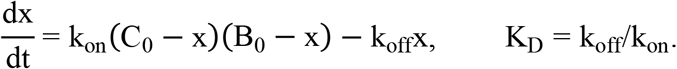

This provides a transparent recognition hypothesis test, not a replacement for spatially resolved binding when the solution is poorly mixed. Both nominal coating concentration and soluble ligand concentration must be controlled: accessible coating density is not determined by nominal coating concentration alone.

### The plate-coated human/goat IgG experiment: why it is an A test conditioned on R

The motivating measurements concern monoclonal anti-human IgG detector antibody interacting with immobilized human and goat IgG in the same 96-well reference reactor. Mouse IgG is close to the original blank. The target human signal is generally stronger than the cross-reactive goat signal. At high coated-IgG and detector-antibody concentrations, particularly after short 10-min incubation, human/goat signals may nearly coincide; dilution and longer 30-/90-min incubations reveal a stronger goat deficit. PBST enhances human/goat separation while retaining much of the high human response; casein decreases nonspecific background but can also depress the high human signal. These are qualitative observations, not raw fitted concentration-by-time curves. The scientific hypothesis is that conventional statements such as “the antibody is highly specific” conceal an assay- and time-dependent recognition output. Here the plate is held fixed to interrogate A. The soluble detection antibody binds the plate-coated species, with accessible coating density, detector-antibody supply, incubation and a common wash all explicitly specified; the actual measured molecular output remains AR (A, R_0_).

### A finite-bath binding solution separates on-rate, off-rate and reagent supply

In the simplest spatially uniform limit, let x_i_ (t) be the molar-equivalent concentration of immobilized complexes of species i ∈ {H, G, M, NS}, B_0,i_ its accessible coating capacity in the same concentration-equivalent unit, and C_0_ the soluble secondary-antibody concentration. The reversible finite-bath model for a separately coated well is

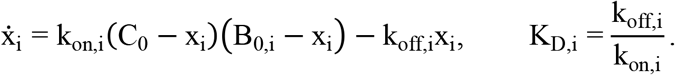

All concentrations in Eq. [eq:Afinite] are in mol/L; converting C_0_, B_0_, x into nM requires k = 10^−9^k_on_ in nM^−1^ s^−1^. Define 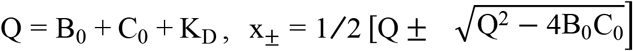 and λ = k_on_ (x_+_ − x_−_) in consistent molar units. The initially unoccupied solution is

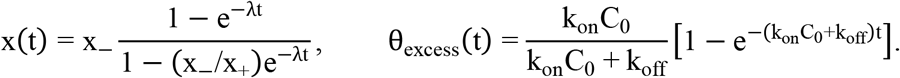

The second expression is a separate pseudo-constant-bath approximation when soluble depletion is negligible. For an illustrative common wash of duration t_w_ with perfect removal and no rebinding, 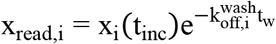. It is a reference limit, not the later finite-volume wash model.

A parsimonious, unmeasured recognition family uses k_on,H_ = k_on,G_ = 10^6^ M^−1^s^−1^, k_off,H_ = 10^−5^ s^−1^ and k_off,G_ = 5 × 10^−4^ s^−1^ (K_D,H_ = 0.010 nM; K_D,G_ = 0.50 nM). It tests whether similar encounter kinetics with substantially different dissociation can create the observed concentration- and time-sensitive selectivity. At the same soluble concentration, site capacity and starting state, the strict inequality k_off,H_ < k_off,G_ gives x_H_ (t) ≥ x_G_ (t) under this minimalist model. Occasional goat-above-human points therefore cannot be assigned to the equal-on-rate model; these require experimental dispersion or an additional physical mechanism rather than an invented rate fit. The predicted time- and concentration-dependent occupancy responses are shown in Figure 2.

**Figure 2.**
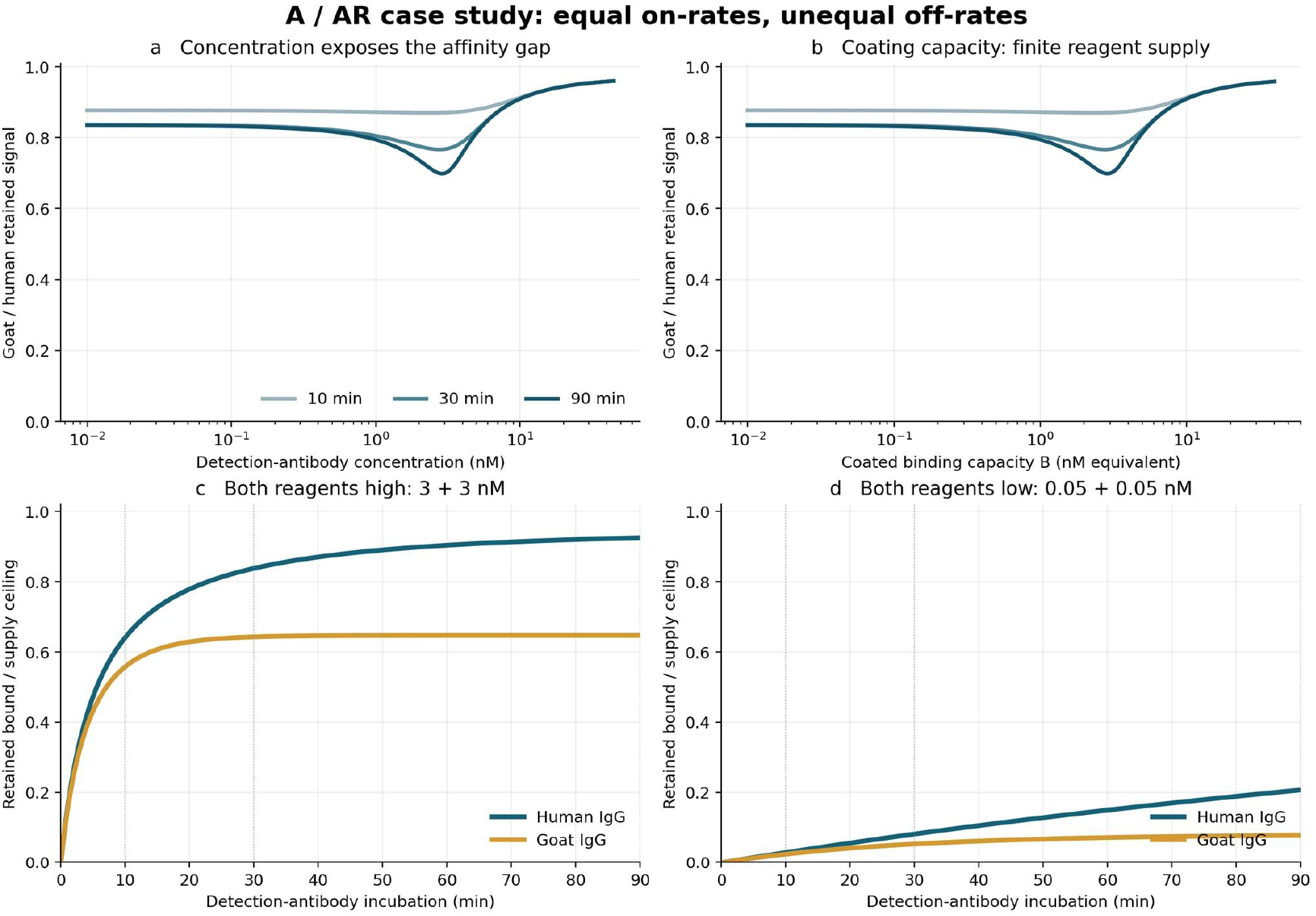
The same nominal target can discriminate differently as time and supply change. Finite-bath model tests high/low coating and soluble detector-antibody concentrations at 10, 30 and 90 min. The response is retained molecular occupancy, not the final optical density. Equal target/off-target k_on_ and unequal k_off_ reveal stronger late-time and low-concentration discrimination.

### Two explicit, nonselective buffer hypotheses: casein slows on-rates; PBST speeds off-rates

The proposed buffer effects are hypotheses about apparent rates under the incubation conditions, not claims that the molecular actions of casein or Tween-20 are proven. Importantly, nonselective means the same multiplicative perturbation across all recognition channels; it does not mean every target and false-target signal changes by the same amount.

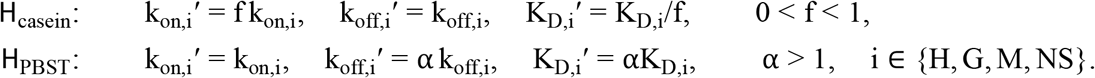

This is a minimal two-scalar intervention: neither buffer is assumed to recognize human, goat or background species separately. The model’s output is nonetheless nonlinear in k_on_C_0_t, C_0_/K_D_, finite available sites, and wash retention. Under PBST, a uniform increase in k_off_ shifts a weak goat interaction into a low-occupancy regime while high-concentration human binding can remain nearly saturated. Under casein, slowing all on-rates penalizes even target binding at finite incubation, while a weak nonspecific channel can fall by a larger fractional amount. Improved target/background contrast is therefore compatible with lower absolute target signal; a better SNR must be calculated from blank variance, not asserted merely from reduced mean NS binding. The modeled effects of these uniform kinetic perturbations on apparent target/background selectivity are illustrated in Figure 3.

**Figure 3.**
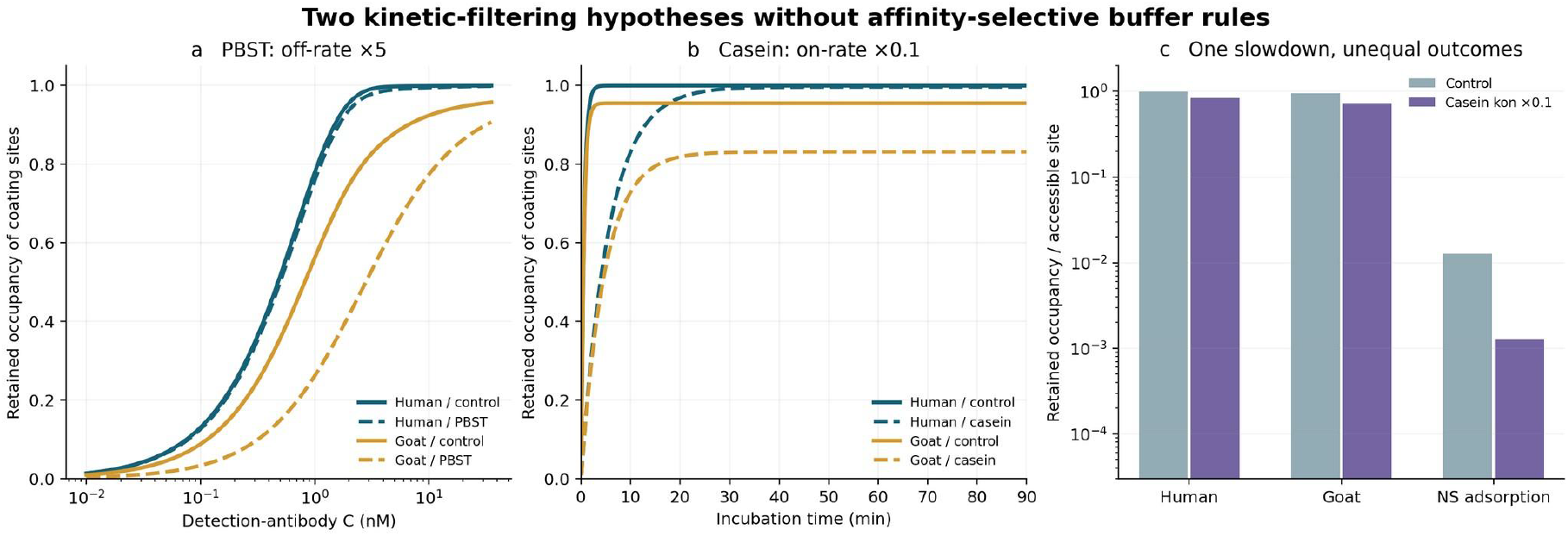
A uniform kinetic perturbation can change apparent target/background selectivity. PBST: an illustrative uniform fivefold k_off_ increase during secondary-antibody incubation; casein: an illustrative uniform tenfold k_on_ decrease. Both are applied equally to target and off-target channels. Occupancy responds unequally because the finite-time binding law is nonlinear. The right panel shows a weak nonspecific-binding channel and is not a measured blank-variance comparison.

### Numerical checkpoints, protocol separation, and tests that distinguish the hypotheses

Under the common equal-on-rate example, the original finite-bath calculations at high coating and high secondary-antibody supply (B_0_ = C_0_ = 3 nM equivalent) predict a retained goat/human ratio of 0.870, 0.767, and 0.699 at 10, 30 and 90 min, respectively; at lower supply (B_0_ = C_0_ = 0.05 nM equivalent) the corresponding ratios are 0.843, 0.654, and 0.377. These numbers illustrate changes in selectivity with time and reagent supply, not direct fitted ELISA optical ratios.

The casein demonstration uses C_0_ = 30 nM and 10 min: a tenfold common reduction of k_on_ changes human pre-wash occupancy from approximately 99.97% to 83.3%, while an independently specified weak nonspecific channel (k_on,NS_ = 10^3^ M^−1^ s^−1^, k_off,NS_ = 10^−3^ s^−1^) falls from approximately 1.34% to 0.14% of its own accessible sites. Those fractions are not directly comparable total molecule counts, and background mean attenuation is not background-SD attenuation. The PBST demonstration holds k_on_ = 10^6^ M^−1^ s^−1^ and multiplies both human and goat incubation off-rates by five; its strong/weak channel separation is a model prediction, not an experimentally measured rate-change factor.The predicted responses to these kinetic perturbations and the corresponding discriminating measurements are summarized in Table 1.

**Table 1.** Predicted effects of uniform kinetic perturbations and discriminating measurements.

| Hypothesis | Expected response at equal R | Discriminating measurement |
| --- | --- | --- |
| Casein: $f k_{on}$ for every channel | Delayed target and false-binding buildup; possible recovery of strong target occupancy at longer time | Full association time series at multiple $C_0$ and coating capacities; compare late equilibrium with early lag |
| PBST: $\alpha k_{off}$ for every channel | Weaker channel more displaced from occupancy saturation; strong human channel relatively protected at high $C_0$ | Dissociation traces after reagent exchange, at matched coverage, and multi-time target/off-target dose responses |
| Shared limitations | Accessible site loss, steric blocking, matrix chemistry and optical interference can mimic either perturbation | Independent label-free rate measurements, controlled coating accessibility and *identical* reporter chemistry/optical readout |

In the human/goat plate experiment, casein and PBST were added to secondary-antibody incubation buffer. Their modeled perturbations must therefore be applied during incubation. The following wash is a common wash protocol, not an additional incubation period in which the hypothetical PBST scaling is automatically re-applied. Wash-stage 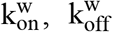 local diffusion and rebinding are a separate AR problem. It is possible for both buffers to affect effective site accessibility or additional chemical channels; the present simple hypotheses are deliberately falsifiable, not exhaustive explanations. Endpoint OD alone cannot uniquely identify intrinsic on/off-rates without independently characterizing available sites, surface avidity, detection-label stoichiometry and downstream T m/T s.

### Recognition–reactor coupling (AR): why affinity alone cannot predict capture

Central test: Change recognition kinetics while holding KD fixed, or change reactor geometry while holding the molecular inputs fixed. If the captured state changes, no isolated affinity score or generic reactor gain can replace the coupled AR calculation. The results are organized from controlled kinetic interventions to geometry, then to assay-specific design decisions.

### Reaction–diffusion boundary problem: the actual physical definition

For species j in a liquid volume Ω_R_ bounded by a reactive surface ∂Ω_R,act_, the benchmark model is

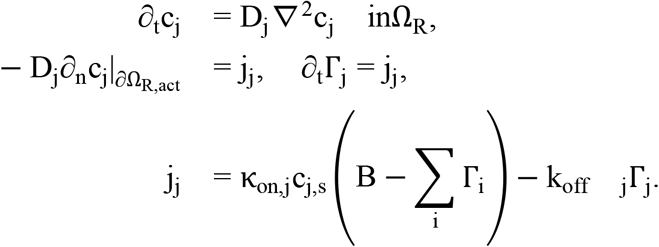

Here κ_on,j_ is the association coefficient expressed in the particle-number-density units of the PDE (volume per molecule per time); molar k_on_ must first be converted using Avogadro’s constant and the chosen spatial units. Here the normal and flux signs follow the convention of the reference formulation; c_j,s_ is the reactor-determined near-wall concentration, B the accessible finite site density, and Γ_j_ the occupied surface density. The coupled diffusion and occupancy fields enforce depletion, finite capacity, competition and rebinding. The wall concentration contains transport memory, schematically 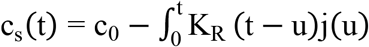 du for appropriate geometry and boundary conditions. The kernel is a Green-function-dependent object, not a universal mean diffusion time.

Intrinsic k_on_ and k_off_ can be measured in a separate assay; the surface complex population cannot be calculated without R. This is the precise meaning of “A and R are not independent outputs.” It does not deny intrinsic affinities. A practical nonseparability test is the response-scale interaction Δ_AR_z = [z (a_2_, R_2_) − z (a_1_, R_2_)] − [ z (a_2_, R_1_) − z (a_1_, R_1_)]: if the benefit of the same kinetic change differs across reactors, this contrast is nonzero. It is a finite-time, output-specific intervention contrast, not a claim that k_on_ itself changes when the reactor is changed. In the dilute initially irreversible limit, two resistances illustrate the same coupling:

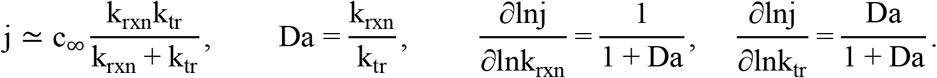

In this limited early-time flux problem, Da ≪ 1 defines a recognition-limited regime: improving the intrinsic surface reaction rate has approximately unit logarithmic leverage while improving transport has little immediate effect. Conversely Da ≫ 1 defines transport-limited flux, for which improving mass transfer has approximately unit leverage while faster binding gives diminishing returns; Da ≃ 1 is the coupled crossover. Equation [eq:series] is an early-time physical intuition only: it omits finite supply, reversibility, wash and history. It must not replace Eq. [eq:pde] in the claimed predictive calculations, and its flux crossover is not automatically the same as the downstream AR-threshold or final ART-LOD crossover.

### Finite-bath reduction, transport clock and exact Riccati trajectory

For comparative scans, express initial molecular inventory per accessible area as A_0_ = C_0_N AV_R_/S_R_ and capture capacity as B (both in molecules/area). Define Γ = X/ S_R_, a = k_on_S_R_/ (N AV_R_) in consistent units, and a reactor-specific effective diffusion factor ϕ_R_ (t). The reduced model is

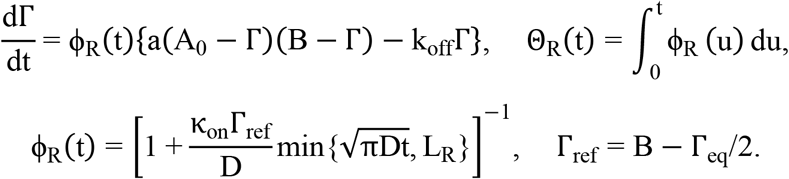

The diffusion scale L_R_ is H/3 for the reflecting plate slab, r/4 for the long cylindrical capillary and a bead-radius-scale local resistance for dilute spheres. These reference closures are geometry dependent, not universal formulae for turbulent mixing or densely packed beads. Crucially, the same factor ϕ_R_ slows the net reversible reaction rather than penalizing k_on_ alone; otherwise the limiting equilibrium would be artificially altered.

Let Γ_−_ < Γ_+_ be the roots of a (A_0_ − Γ) (B − Γ) − k_off_Γ = 0. Explicitly,

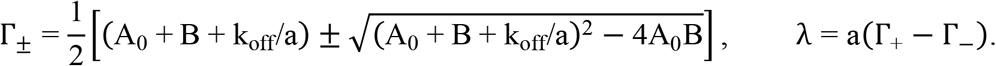

For initially vacant sites, integration of the Riccati equation in effective time yields

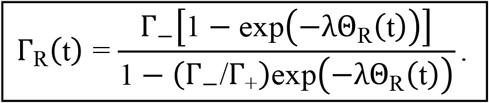

This is an exact solution of the reduced ODE, not an exact solution of the spatial PDE. It preserves finite analyte supply and correct closed-bath equilibrium but cannot reproduce arbitrary spatial memory, mixing, multivalent kinetics or wash exchange. The implementation reproduced 108 archived predictions at K_D_ = 0.1 and 1 nM to floating-point precision; those checks show software consistency, not physical validation. Independent finite-volume PDE comparisons cover only a subset of plate/TOI parameter space and report potentially large errors in extreme fast-k_on_ plate conditions. Bead extremes and k_on_ ≥ 10^7^ M^−1^s^−1^ remain exploratory extrapolations.

### Reactor geometry and interpretable observables

The controlled comparison uses D = 4.46 × 10^⁻11^ m^2^/s, B = 1,000 accessible sites/ μm^2^, and C_0_ = 1 nM for time courses. The geometric and transport characteristics, together with the corresponding sample volumes, reactive surfaces, and analyte inventories, are summarized in Table 2.

**Table 2.** Geometric and transport characteristics of the three reference reactor configurations.

|  | **96-well**<br>**plate** | **TOI capillary** | **10k beads** |
| --- | --- | --- | --- |
| Sample volume | 100 $\mu\text{L}$ | 8.836 $\mu\text{L}$ | 50 $\mu\text{L}$ |
| Reactive surface | 32 mm<sup>2</sup><br>bottom | 0.75-mm ID<br>×20-mm wall | 10,000 beads; 3-$\mu\text{m}$<br>diameter |
| R transport scale | slab $H/3$ | cylinder $r/4$ | dilute-sphere local<br>model |
| At 1 nM, $A_0$<br>(molecules/$\mu\text{m}^2$) | 1882 | 112.9 | geometry-dependent |

Three observables must be kept distinct: surface density Γ_R_ (C, t), number captured X_R_ = S_R_Γ_R_, and recovery η_R_ = X_R_/ (CN AV_R_). None alone is a measured signal or LOD. This distinction explains why a small-volume reactor may have high recovery but modest absolute surface density or lower total captured molecules.

### Result 1: identical affinity, different time-dependent capture

At fixed K_D_ ∈ {0.1,1,10} nM, six paired rates k_on_ = 10^3^, …, 10^8^ M^−1^s^−1^ with k_off_ = K_D_k_on_ are transported into each reactor. Thus each nominal affinity labels six different transient curves. At K_D_ = 0.1 nM, increasing k_on_ from 10^5^ to 10^6^ M^−1^s^−1^ with proportionate k_off_ increase gives a 10-min capture-density gain of 2.86 × (plate), 2.16 × (TOI) and 7.42 × (beads); at 240 min those ratios fall to 1.36 ×, 1.000 × and 1.26 ×. Same A change; distinct benefits because of R and endpoint time.

### Complete transient trajectories, not only an endpoint snapshot

The main-text exemplar fixes K_D_ = 1 nM while varying six kinetic pairs in each of three reactors; it preserves both early and full incubation windows. Full K_D_ = 0.1 and 10 nM counterparts remain available in Supplementary Figs. S1 and S2. Endpoint maps are a different observable, not interchangeable with time traces. These curves are not redundant with the following 15-min summary: they show where the apparent kinetic benefit grows, crosses other traces, or disappears as reactor-specific inventory and equilibration limits become dominant. Each panel uses the identical named antibody pair across three different reactors; changing R is never represented as merely rescaling the same capture history. The full time-dependent capture trajectories for the matched kinetic pairs across the three reactor configurations are shown in Figure 4.

**Figure 4.**
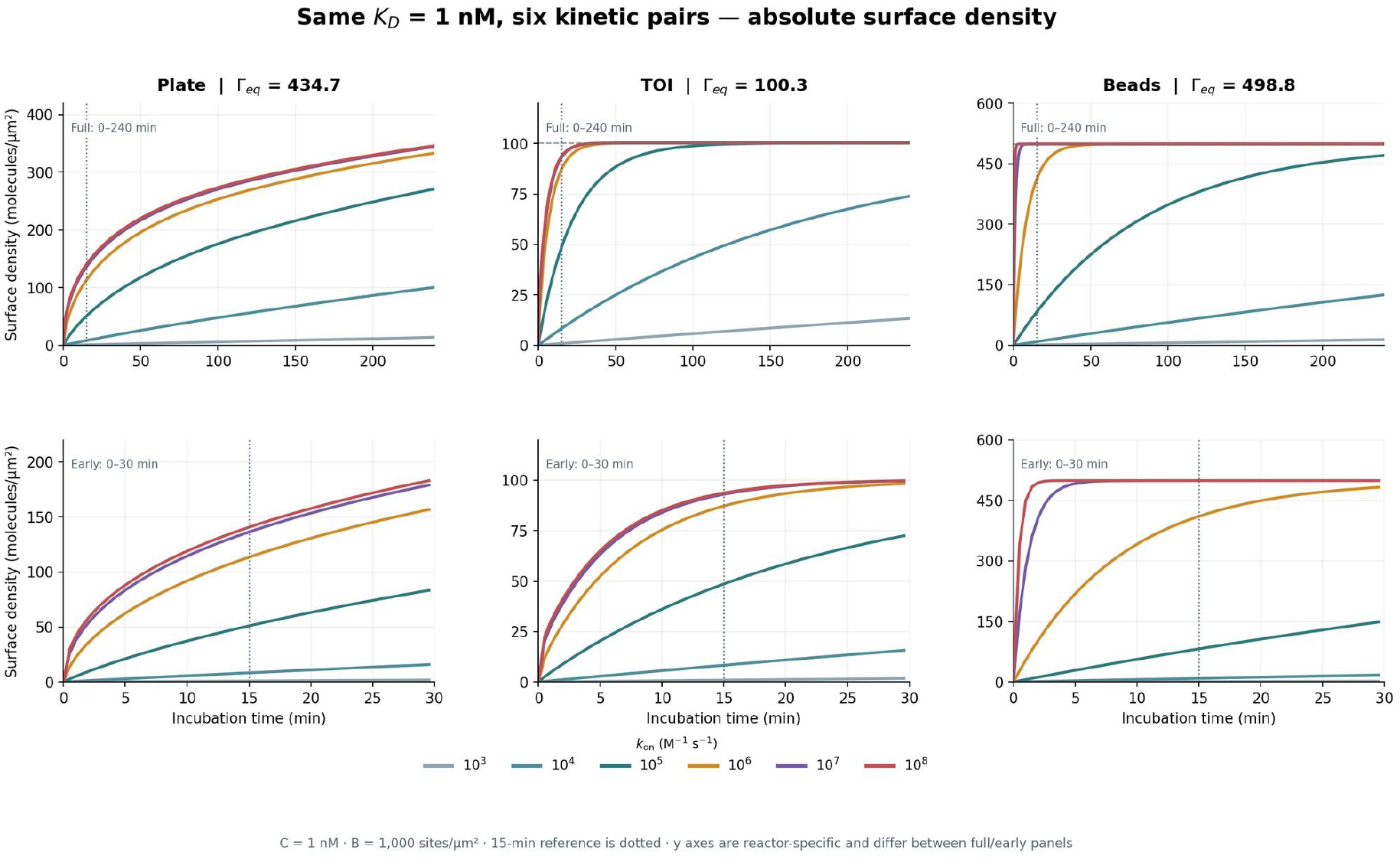
A single K_D_ supports many distinct time trajectories in three reactors. At K_D_ = 1 nM, six matched (k_on_, k_off_) pairs and three reactors are shown at full and early incubation times. Companion K_D_ = 0.1 and 10 nM kinetic families are preserved in Supplementary Figs. S1 and S2. These are simulations, not experimental binding curves.

### An endpoint comparison is a cross-section of those trajectories

The matched k_on_ = 10^6^ 15-min numerical checkpoints are plate / TOI / beads = 148.6/93.5/560.2 molecules/µm^2^ at K_D_ = 0.1 nM; 113.3/87.2/410.4 at 1 nM; and 63.2/50.6/90.8 at 10 nM. Changing K_D_ from 10 to 0.1 nM at fixed k_on_ therefore yields 2.35 ×, 1.85 × and 6.17 × density increases in the three reactors. These are conditional predictions at 15 minutes, not platform-invariant antibody quality factors.

### Rate interventions provide an independent kinetic test

A second controlled intervention fixes k_on_ and changes only k_off_ to produce K_D_ = 0.1, 1, and 10 nM. The supporting calculation (Supplementary Figs. S4–S5) covers two fixed association rates: 10^5^ and 10^6^ M^−1^s^−1^. The corresponding six families of long/early time trajectories are shown below. They are not interchangeable with Experiment III, where affinity is held fixed and both rates are varied simultaneously. At short time, changing dissociation can have modest influence when rebinding or association kinetics dominate; at later time, the equilibrium loss becomes more visible. The temporal response is reactor-dependent throughout.

### Result 2: controlled on/off perturbations change dose–capture maps

The important assay-facing AR output is Γ_R_ (C, t) across concentration, not a single 1 - nM time course. The calculations sweep C from 0.01 pM to 100 nM at 15 minutes, with three carefully controlled interventions. Figures 5–7 show the respective response maps:

**Figure 5.**
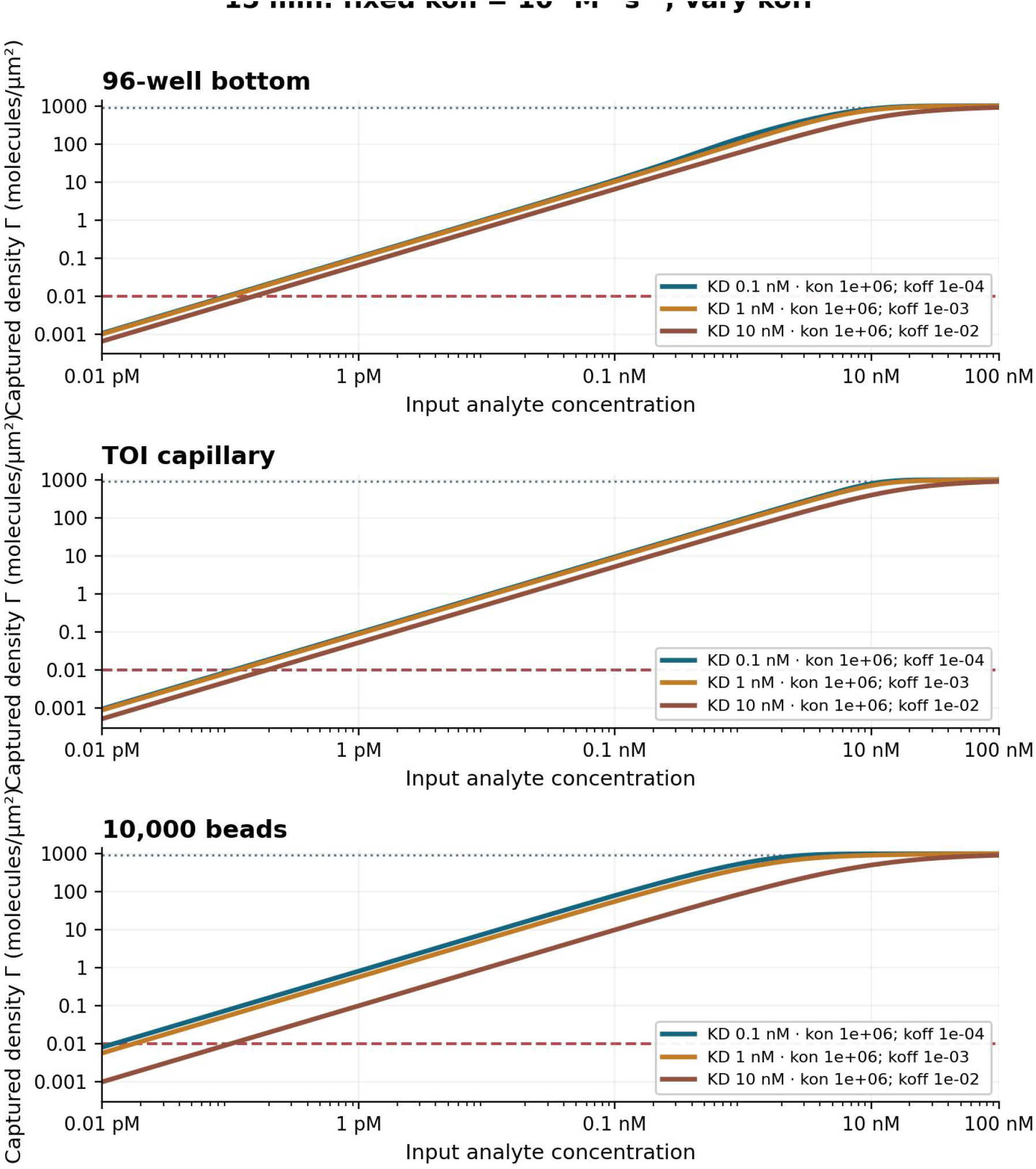
Changing k_off_ at fixed k_on_ changes the finite-time capture curve. Concentration-to-capture-density curves for plate, TOI and beads with k_off_-controlled K_D_ differences. Same concentration scale and absolute surface-density units; dashed and dotted horizontals mark Γ_L_ and 0.9B, not instrument thresholds.

**Figure 6.**
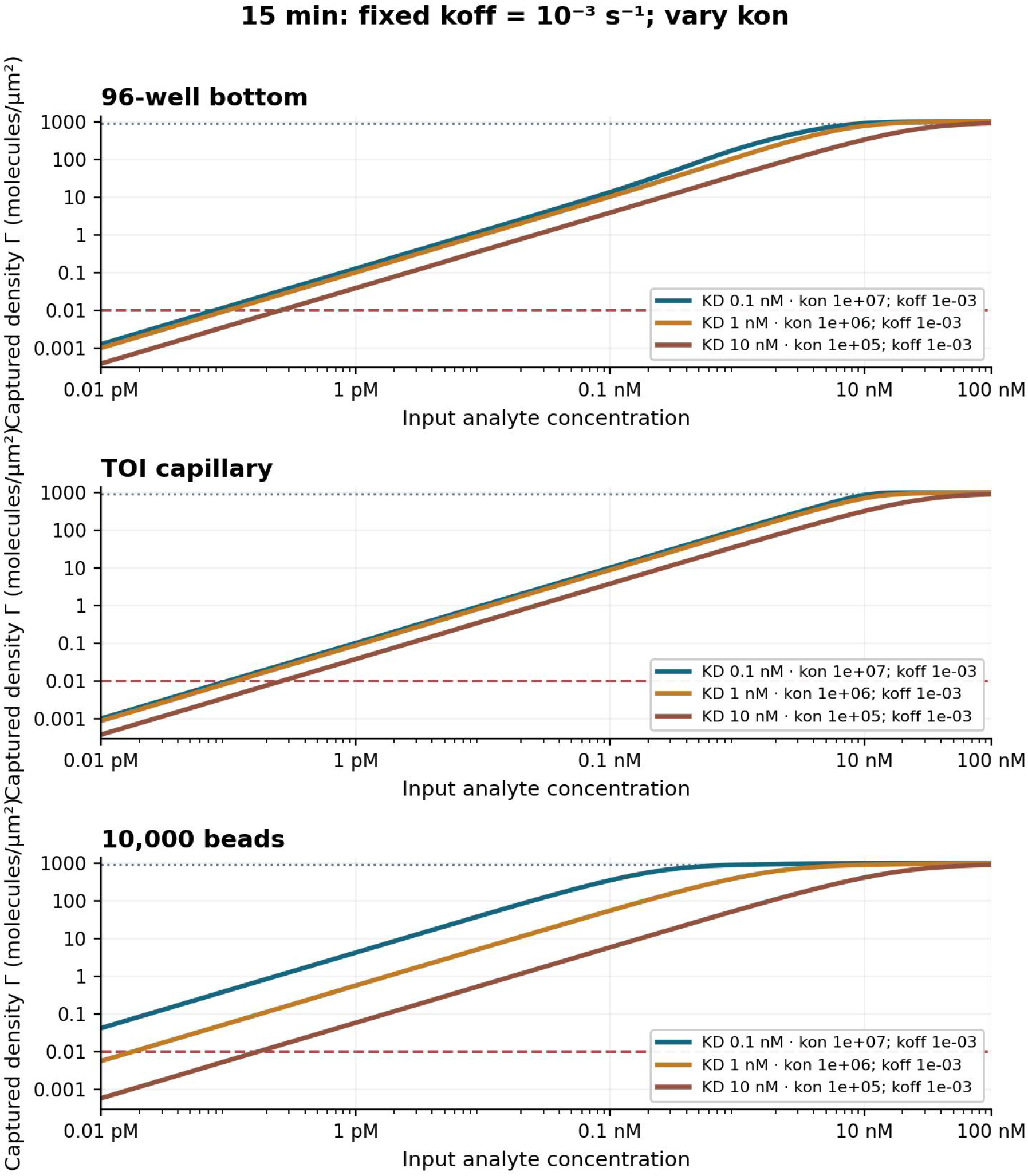
Changing k_on_ at fixed k_off_ is not equivalent to changing k_off_. Identical affinity labels obtained through changes in k_on_ alone produce different finite-time concentration-response functions from the preceding fixed-on-rate results. This contrast is the empirical target for the statement that K_D_ does not uniquely determine a fixed-time immobilized dose response.

**Figure 7.**
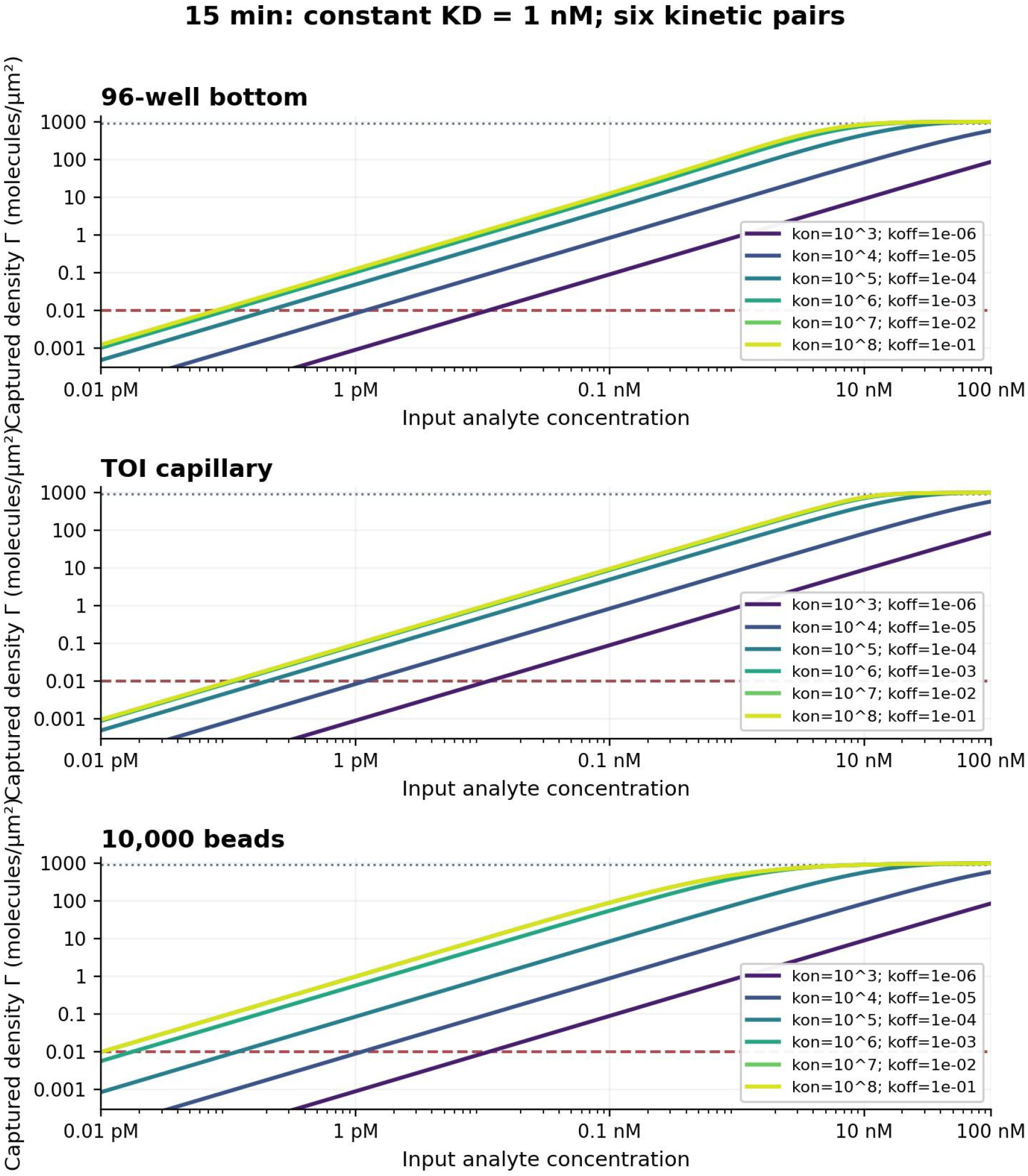
Equal K_D_ does not produce one concentration-response function. All six matched kinetic pairs at K_D_ = 1 nM are evaluated at the same 15-min time point across plate, tube and bead reactors. The corresponding full K_D_ = 0.1 and 10 nM concentration families are in Supplementary Figs. S6 and S7. These are model outputs, not experimental data.

1. Fixed on-rate: k_on_ = 10^6^ while k_off_ = 10^−4^, 10^−3^, 10^−2^ s^−1^ generates K_D_ = 0.1,1,10 nM.
2. Fixed off-rate: k_off_ = 10^−3^ s^−1^ while k_on_ = 10^7^, 10^6^, 10^5^ M^−1^ s^−1^ generates the same affinity labels by a different physical intervention.
3. Fixed affinity: for each of K_D_ = 0.1,1,10 nM, six coupled k_on_, k_off_ pairs test kinetic diversity without changing equilibrium K_D_.

The absolute-density lower reference Γ_L_ = 0.01 molecules/ µ m^2^ and upper 90%-site threshold Γ_U_ = 900 molecules/µm^2^ define an AR-only window, D_AR,90_ = log_10_ (C_U,90_/C_L_) if the upper crossing exists within the scanned input range. These are not the final analytical LOD, LOQ or T s roof. Right-censored C_U,90_ values must not be fabricated from the 100-nM plotting boundary.

Experiment III is a third distinct physical intervention: at each fixed K_D_ ∈ {0.1,1,10} nM, six on/off pairs are varied together. The complete three-reactor concentration-to-surface-density families must all remain visible, not reduced to one representative affinity curve. The x-axis is analyte concentration rather than incubation time; the same affinity can exhibit distinct effective slopes and threshold crossings under a finite-time analytical protocol.

### Response trajectories and AR windows measure different properties

The complementary experiment with fixed k_off_ = 10^−3^ s^−1^ changes k_on_ alone. Even at C_0_ = 1 nM, its 0–240-min time curves differ from the fixed-on-rate experiment because the rate-limiting physics is changed, not merely the equilibrium parameter label. The AR-only density-threshold concentration span (defined by Γ_L_ and Γ_U_) is therefore a property of a specified A, R, t setting, not an intrinsic readout or assay dynamic range.

### Washing remains part of coupled AR physics

Washing can release bound species, change near-wall concentration, and permit local rebinding. In the finite-exchange-layer model, ∂_t_c_w_ = D_w_∂_xx_c_w_ on 0 < × < δ with c_w_ (δ, t) = 0, ∂_t_Γ_w_ = κ_on,w_c_w_ (0, t) (B − Γ_w_) − k_off w_Γ_w_, and the surface diffusive flux matched to this net reaction rate. The initial Γ_w_ 0 is the post-incubation surface state, not an equilibrium reset. In a perfect sink with no rebinding, the retained fraction is exp (− k_off_t_w_); with finite exchange, reversible transport must be solved again. For t_w_ = 30 s and k_on_ = 10^6^, the ideal sink loss bound is 0.30%, 2.96% and 25.92% for K_D_ = 0.1,1,10 nM (corresponding k_off_ = 10^−4^, 10^−3^, 10^−2^ s^−1^). At higher wash-induced off-rate stress or longer exchange residence, wash can dominate the surviving reporter-capable population. The rebinding analysis explicitly varies a not-yet-calibrated effective exchange distance δ, and must not be mistaken for measured 0.05%-PBST hydrodynamics.

### Result 3: a physically audited reactor model reveals design trade-offs

Historical contribution and model refinement. The TOI platform reported in hLife establishes a concrete microvolume immunoassay setting in which reactor design matters experimentally.[1] In a complementary theoretical paper published earlier in 2025, Lyu et al. made the encounter-time problem quantitative using a first-passage-time (FPT) construction, systematically relating tube radius, incubation duration, collision/immobilization probability P_on_ and captured density.[2] Their model and experiments motivated examination of a nonmonotonic radius optimum; the quoted best radii at 600 and 3600 s were 0.27 and 0.66 mm, respectively, within that approximation. That study also reported controlled experimental tests of the overall model trends. Its conceptual contribution remains: neither antibody affinity nor miniaturization alone defines a successful short-time assay, and reactor dimensions should be selected by the stated objective and operating time. Here ART preserves that design question while checking whether the single-exponential first-passage closure gives the correct local-density trend in a spatially resolved, mass-conserving reversible-binding cylinder. The present reference sweep holds tube length fixed rather than holding reactor volume fixed; its design window should therefore not be interpreted as an exact reproduction of the published FPT sweep.

### One experimentally legible design intervention: fixed *l* = 2 mm

Fix *l*_0_ = 20 mm, incubation t_i_ = 900 s, no imposed axial flow, accessible capture density B = 1000 sites/µm^2^, D = 44.6 µm^2^/s and k_on_ = 10^6^ M^−1^s^−1^. Test K_D_ = 0.1 and 1 nM by setting k_off_ = k_on_K_D_. For each radius r, the reactor volume and total initial target inventory are not held fixed:

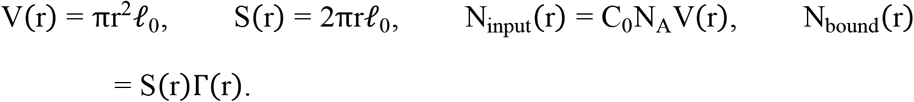

The maximum possible bound density follows by placing all input molecules on the accessible sidewall:

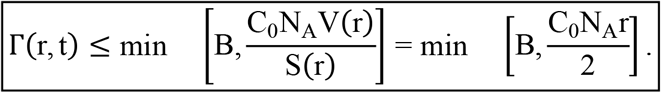

Use C_0_N_A_ consistently as molecular number density. The inventory ceiling rises linearly with radius; at sufficiently small radius, even perfect capture cannot yield a high per-area output. Fixing length does not eliminate that small-radius penalty. Moreover, N_input_ ∝ r^2^ and N_bound_ ∝ rΓ, so the same change in radius affects total capture and surface density differently.

### Model 1: nonlinear conservative radial transport and reversible wall binding

The full axisymmetric reference solves

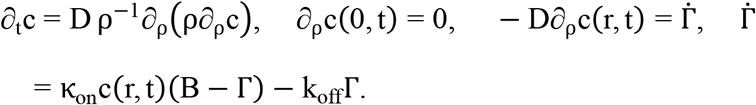

We start from uniformly dissolved molecules and an unoccupied wall. Axial uniformity and reflecting ends reduce the 20-mm tube to a radial finite-volume model; the tube length multiplies volume and surface equally in the mass balance, so Γ r itself is independent of *l*_0_ under these particular boundary assumptions. The nonlinear PDE retains site depletion, finite molecule inventory and dissociation. Incubation and an explicitly separate 30 -s ideal-sink wash are not conflated; plots of incubation-end density use Γ (t_i_), while threshold curves use 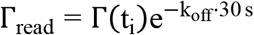 without wash rebinding.

### A falsifiable approximation: the first-passage exponential closure

For uniformly initialized independent particles with a Robin sidewall, the mean first-passage time and the approximate variance used in the archived surrogate figures are

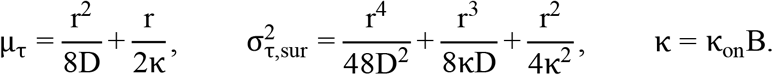

For the same uniform initial condition and Robin sidewall, the independent backward-moment equations yield the exact first-passage variance

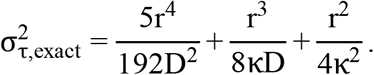

The archived comparison used the smaller r^4^/ (48D^2^) coefficient, which is an approximation rather than the exact cylinder variance. For traceability we retain those numerical surrogate curves unchanged instead of claiming that they were recomputed with the corrected variance. Their peak locations and threshold values should be interpreted only within that archived approximation; the qualitative asymptotic problem with global single-exponential survival remains.

Using a moment-informed, single-exponential survival surrogate motivated by the FPT treatment of Lyu et al., set

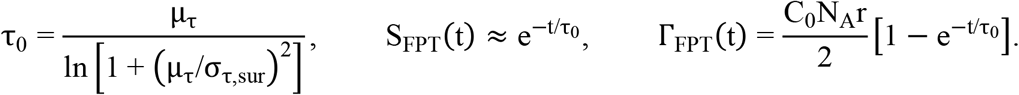

This one-parameter exponential is informed by the first two FPT moments but does not match both moments exactly (an exponential distribution has equal mean and standard deviation). It is a comparison surrogate, not an equivalent solution of Eq. [eq:toi-fixedlength-pde]: the published P_on_ parameter is a collision/immobilization probability and is not experimentally interchangeable with k_on_. Here we match a Robin wall conductance κ to the unsaturated PDE boundary and keep the same D, B, geometry and incubation time. The FPT surrogate omits site occupancy, time-dependent rebinding and in-incubation k_off_; we multiply it by the same ideal-sink 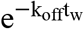 for a declared wash-side comparison. Accordingly, comparisons are mechanism audits, not exact reproduction of the published P_on_ figures.

### Result: an FPT radius peak is not a model-independent density optimum

Figure 8 provides the principal model-discrimination result and reports the full nonlinear Γ (r, C_0_) over C_0_ = 0.001–100 nM for both affinity values after 15 min. At C_0_ = 1 nM and K_D_ = 0.1 nM, the PDE predicts approximately Γ = 30.5, 72.8, 90.6, 103.3, 109.7, and 115.3 molecules/µm^2^ at r ≃ 0.10, 0.26, 0.374, 0.61, 0.99, and 2.50 mm. The surface density increases toward a large-radius plateau: it does not exhibit a finite-radius peak within the modeled domain. At the same reference point, the FPT exponential approximation instead predicts a peak near r = 0.37 mm and a subsequent decline; its r = 2.5 mm prediction is about 20.7 molecules/µm^2^ compared with the PDE value of 115.3. The visual distinction is a numerical finding under the stated, archived exponential approximation; its exact numerical peak would require recomputation with the independent variance correction described above.

**Figure 8.**
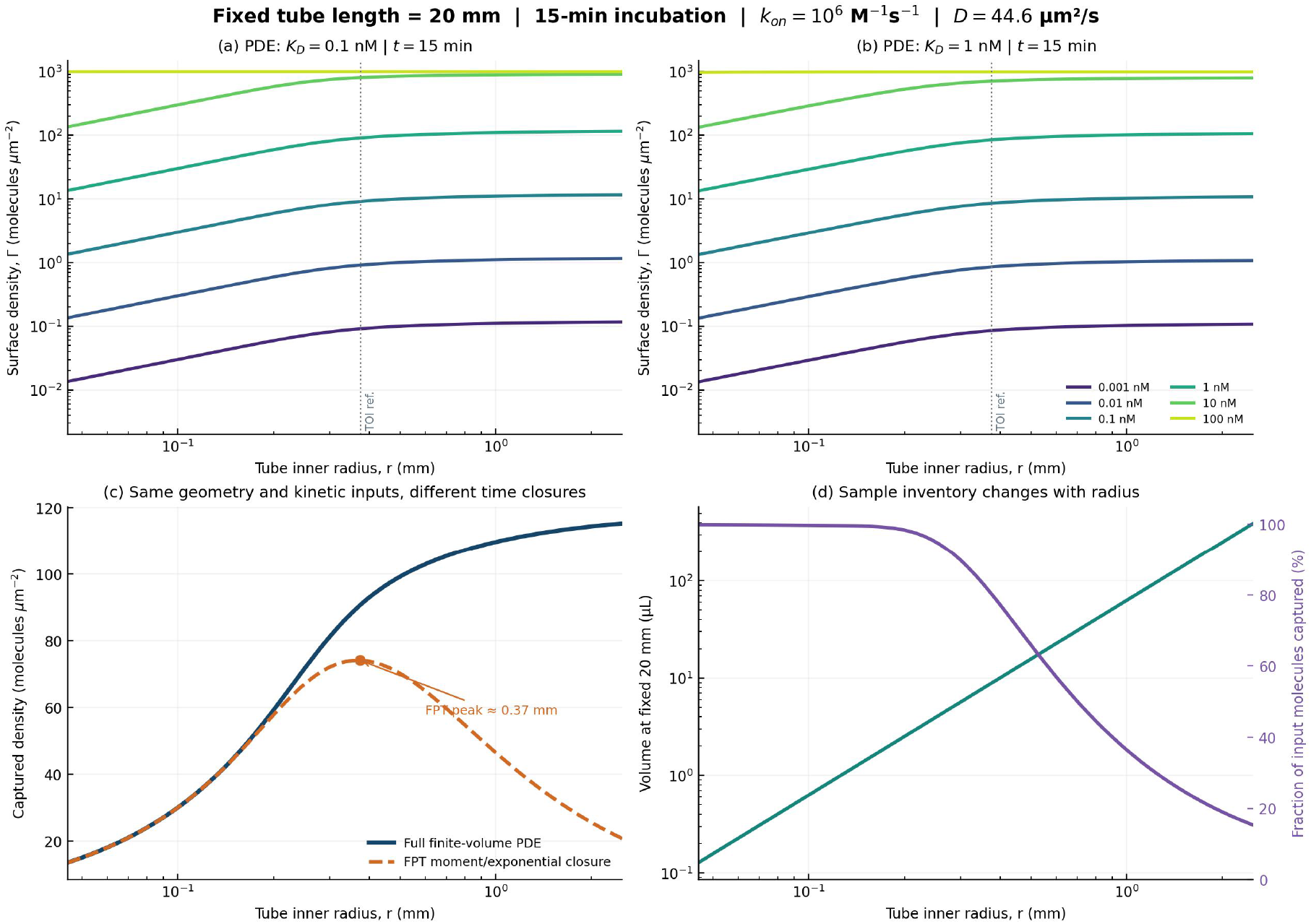
Spatially resolved AR refines the FPT-derived radius-density prediction. (a,b) Full nonlinear reversible radial-PDE prediction with six molecular input concentrations; horizontal axes are inner radius (mm), vertical axes are surface capture density (molecules/μm^2^), analogous to the axes in Lyu et al. (c) Explicit comparison of the same kinetic inputs in a spatial PDE and moment-informed exponential FPT closure at 1 nM, K_D_ = 0.1 nM; only the latter produces an interior radius maximum. (d) The reagent volume increases as r^2^ at fixed 20-mm length while the captured fraction drops with radius. All curves are theoretical and do not include optical noise or calibrated signal conversion.

The difference has a precise asymptotic explanation. When 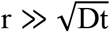, particles initially within a diffusion length of the wall still contribute during the assay. For a perfectly absorbing wall the short-time local planar limit is

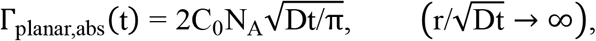

which is nonzero and independent of radius. A finite reactive velocity or a finite dissociation rate modifies its value, not the fact that the local near-wall population survives the large-radius limit. Conversely, the single-exponential MFPT closure has τ_0_ ~ r^2^/D and therefore Γ_FPT_ ~ (C_0_N_A_r/2) t/τ_0_ ∝ 1/r at fixed time. A global mean first-passage time is thus an insufficient proxy for the early-arrival tail from near-wall starting positions. This difference is most pronounced at large radius and short reaction time; it does not invalidate the utility of MFPT calculations for first-passage statistics, but constrains applying a one-parameter exponential survival law to entire endpoint concentration curves.

### An Lyu-style surface-density threshold, not a true assay LOD

The source article converted its modeled signal into a theoretical LLOD by intersecting Γ (C_0_) with a specified density of 0.01 particles/µm^2^. We reproduce that decision rule as an explicitly conditional AR metric:

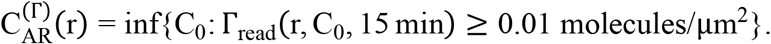

This neither fixes the blank variance nor specifies the chemiluminescent T_m_ and optical T_s_ readout, so it is not a measured or complete ART LOD. The 30-s ideal-sink wash is retained for the ART scenario and differs from the source paper’s P_off_ = 0 simulation condition. Supplementary Fig. S12 shows that the FPT surrogate produces a model-dependent finite-radius minimum in this input-concentration threshold under the present parameter mapping, whereas the conservative PDE declines towards a transport-limited plateau. At the 0.374-mm reference radius, the modeled K_D_ = 0.1 - nM threshold is about 110 fM by PDE and 135 fM by the FPT surrogate; at 2.5 mm the comparison is 86 fM versus 483 fM, respectively. These specific values are hypothetical AR thresholds based on the 0.01/µm^2^ rule.

### Independent exact first-passage check: no exponential-survival shortcut

For a perfectly absorbing circular wall at ρ = r, a uniformly initialized finite cylinder with reflecting ends has an independent analytical solution. Let α_n_ be the positive zeros of J_0_. The exact survival fraction and captured wall density are

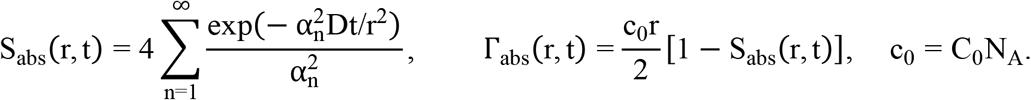

This solution uses the complete first-passage-time distribution, not an exponential ansatz built from its mean. In the finite-κ, irreversible Robin case, the corresponding analytical expansion has eigenvalues defined by λ_n_J_1_ (λ_n_) = Bi J_0_ (λ_n_), Bi = κr/D, and weights 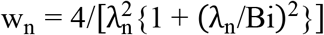; consequently 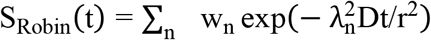Both eigenexpansions reduce at large r to a finite near-wall capture flux. The planar Robin limit can be written analytically:

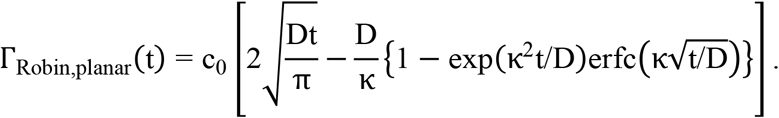

Thus neither the ideal absorbing nor the finite-reactivity problem predicts density decaying to zero merely because the tube radius becomes large. The MFPT-exponential closure instead uses a characteristic time growing as r^2^/D, and spuriously sends the entire population’s capture fraction to zero at a rate that suppresses the locally arriving near-wall subpopulation. Independent absorbing-wall Bessel and partially absorbing Robin solutions verify the short-time near-wall contribution at 30, 60, 300 and 900 s (Supplementary Fig. S24), without using the nonlinear numerical PDE as its own validation. Importantly, the corrected wall-density response does not eliminate a useful reactor comfort zone. Figure 9 combines the historical FPT peak, the updated PDE result and an explicitly conditional density–recovery design utility. The FPT study remains the citable precursor that first formalized the TOI size/encounter-time question. The narrower finding here is that the particular single-exponential closure under the present matched, fixed-length passive-reactor assumptions is not interchangeable with the spatially resolved early-arrival distribution for wall-density predictions. This is not a refutation of FPT theory as a whole or of the original study’s experimental observations.

**Figure 9.**
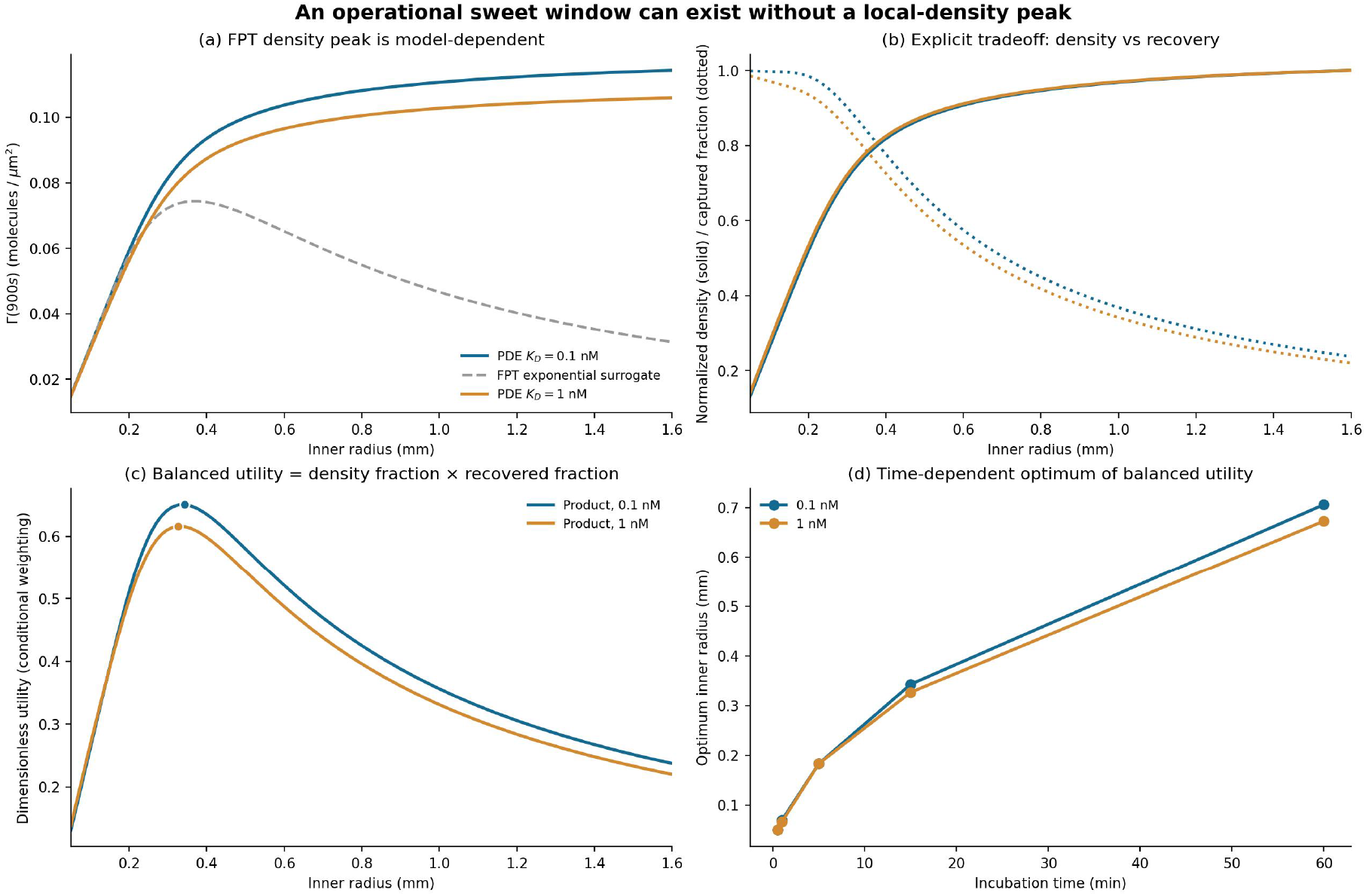
A conditional TOI operating window survives correction of the FPT wall-density optimum.(a) The historical single-exponential FPT surrogate produces an intermediate-radius wall-density peak, whereas the mass-conserving PDE predicts a monotonic-to-plateau local-density response under the stated fixed-length conditions. (b) Smaller radii favor recovered fraction while larger radii favor local density. (c) A clearly defined, dimensionless illustrative utility (normalized density multiplied by recovered fraction) admits an interior maximum without claiming that a measured LOD does. (d) That optimum varies with incubation time; the approximate 0.25–0.50-mm range is a candidate operating window for the representative conditions, not a universal tube-size prescription. The chosen multiplicative utility and its equal weights are design assumptions, not a derived detection law.

### Does lower concentration or shorter incubation restore a radius optimum?

To test the design hypothesis directly, we recalculated the nonlinear reversible finite-volume PDE for four incubation times (30, 60, 300, 900 s), four input concentrations (10 fM, 1 pM, 0.1 nM, 1 nM) and two dissociation constants (K_D_ = 0.1 and 1 nM at fixed k_on_ = 10^6^ M^−1^s^−1^). All calculations hold the sealed tube length at 20 mm, the accessible density at 1000 sites/ µ m^2^ and D = 44.6 µ m^2^ /s. Supplementary Fig. S13 gives all 32 combinations (four concentrations × four times × two K_D_ values); every predicted surface-density trace increases from a molecule-inventory-limited small-radius regime toward a large-radius plateau, without an interior declining branch across 0.035–3.5 mm. This is a conditional model result, not a theorem covering all microfluidic systems. At low occupancy, Γ scales almost linearly with C_0_, so reducing concentration alone does not change the ordering of radii; it does, however, make actual molecular shot noise and finite-copy sampling more important.

### What microfluidic confinement does improve: recovery and sample economy

At fixed C_0_ and t, surface density Γ and fractional recovery η = Γ/ (c_0_r/2) need not improve in the same direction. For C_0_ = 1 pM and K_D_ = 0.1 nM, the reference PDE predicts at 60 s approximately Γ = 0.0194 molecules/µm^2^ and η = 64.4% for r = 100 µm, compared with 0.0230 molecules/ µm^2^ and η = 3.05% for r = 2500 µm. After 900 s, the corresponding values are 0.0300 and 99.7% versus 0.1164 and 15.5%. Thus a 25-fold reduction in radius decreases local 60-s density by only about 16% while increasing the recovered fraction by about 21-fold. This does not imply an equally improved analytical LOD, since reaction area, total photons, blank distributions and the optical collection geometry are not held constant. It does demonstrate an unambiguous engineering advantage when analyte volume, assay speed or recovery efficiency is limiting. The contrasting effects of tube radius on surface capture density and fractional recovery are illustrated in Figure 10.

**Figure 10.**
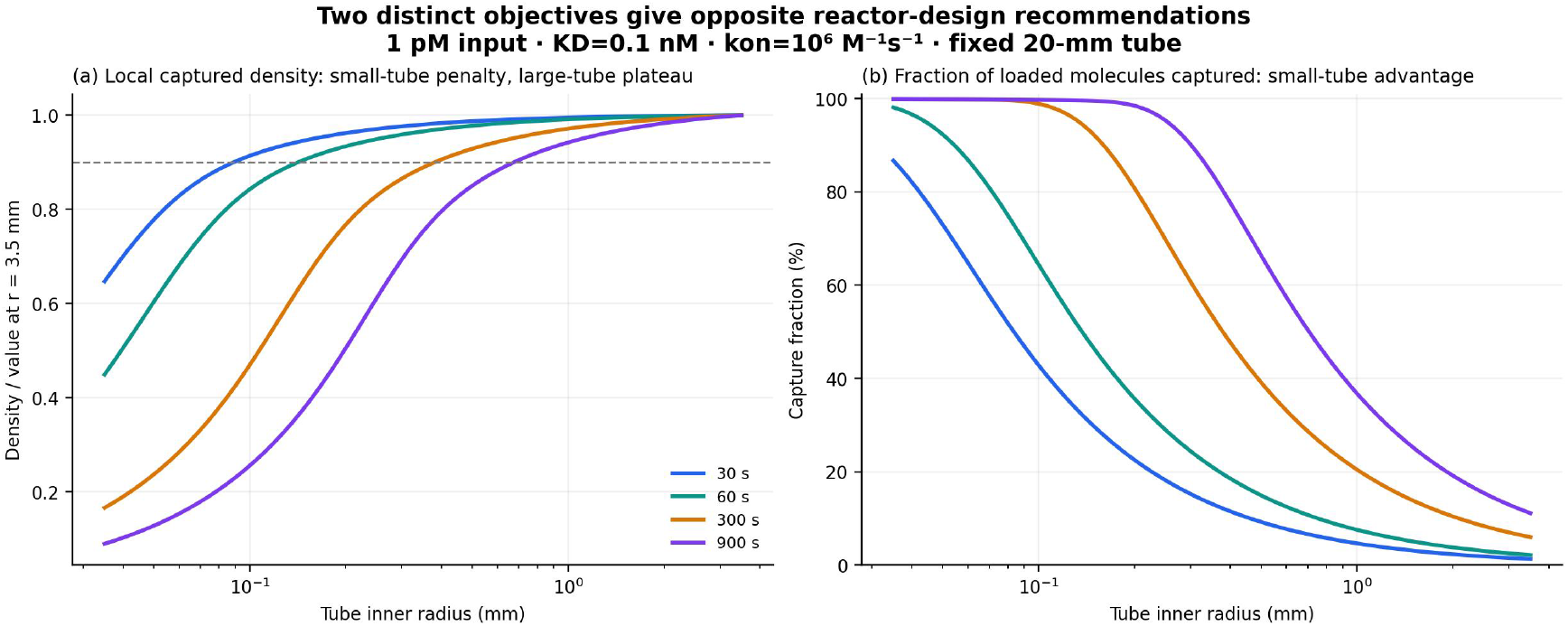
Surface density and sample recovery reward different geometries. Fixed 20-mm tube at 1 pM and K_D_ = 0.1 nM. Left, wall density normalized to the modeled 3.5-mm-radius reference, approaching a finite plateau as radius grows. Right, fraction of input molecules captured, which strongly favors narrow bores under the same fixed-time constraint. The 90% line marks a design knee, not a true maximum or an instrumental LOD boundary.

### An operational design window: density, recovery, absolute capture and time-to-80%-equilibrium

The absence of a finite-radius maximum in local wall density under the static, homogeneous cylindrical model does not invalidate useful size optimization. We distinguish four observables at fixed tube length *l* = 20 mm: the captured wall density Γ (molecules/ µ m^2^), total captured molecule number N_cap_ = 2πr*l*Γ, the recovery fraction η = N_cap_/ (c_0_πr^2^*l*) and an explicitly defined kinetic response time. Neither Γ nor recovery alone is an instrumental LOD, and a higher total capture count also requires accounting for the larger input volume and optical collection geometry.

For finite-bath reversible binding, the relevant saturation target is the actual concentration- and geometry-dependent equilibrium, not 80% occupation of the total coating sites. With the accessible capacity B, initial number concentration c_0_ and finite-bath equilibrium free concentration c_eq_, mass conservation and Langmuir binding give

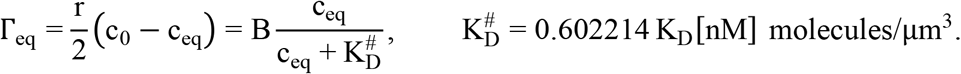

Our revised kinetic metric is

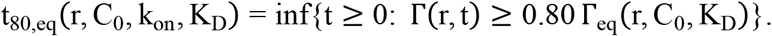

The benchmark is defined consistently as 80% of each condition-specific equilibrium throughout this analysis. It differs from the moment-informed exponential FPT surrogate time t_80,FPT_ = τ_0_ln5: the latter represents approach to a model-specific, irreversible capture-inventory ceiling, not the finite-bath reversible surface equilibrium. Since a weaker antibody can reach 80% of a lower equilibrium density sooner, the relative kinetic metric must always be displayed alongside its actual wall-density and molecule-count outputs.

### A wide affinity spectrum: ordinary 10-nM antibodies must not be omitted

We retained the same mass-conserving radially discretized reversible cylindrical PDE, tube length 20 mm, D = 44.6 µ m^2^ /s, accessible site capacity B = 1000 sites/ µ m^2^, and finite-bath equilibrium definition above. The expanded sweep contains five affinities K_D_ ∈ {0.01,0.1,1,10,100} nM, three association rates k_on_ ∈ {10^5^, 10^6^, 10^7^} M^−1^s^−1^, four analyte inputs (0.001,0.1,1,10) nM, and ten directly simulated inner radii (0.10,0.20,0.25,0.33,0.375,0.40,0.45,0.50,0.70,1.00) mm: 600 parameter combinations. The sweep deliberately spans weak-to-ultrahigh apparent affinities and includes the ordinary 10-nM class; all rate and concentration choices are modeled conditions rather than measured antibody properties. At each fixed k_on_, the five K_D_ values change k_off_ = 10^−9^K_D_ [nM]k_on_; increasing k_on_ at fixed K_D_ changes both on and off rates and is a distinct controlled comparison.The resulting trade-offs among equilibration time, surface capture density, fractional recovery, and absolute captured molecule count are shown in Figure 11.

**Figure 11.**
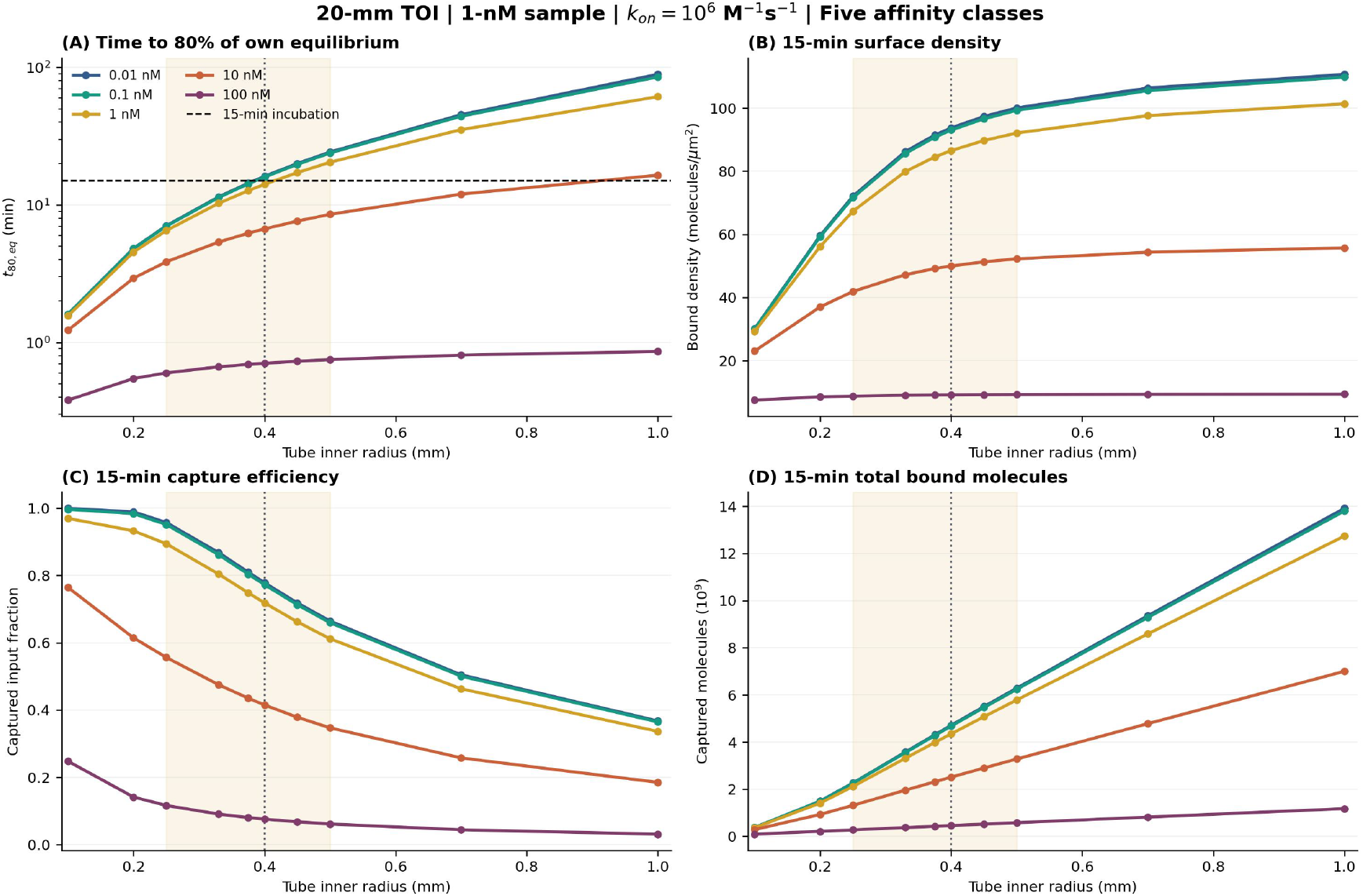
Affinity, incubation speed, density and recovery are distinct objectives. The fixed 20-mm tube contains 1-nM analyte, with k_on_ = 10^6^ M^−1^ s^−1^; colors distinguish K_D_ = 0.01,0.1,1,10,100 nM. Four different measures are plotted against radius: (a) time to 80% of each condition’s own finite-bath equilibrium, (b) 15-min captured density, (c) 15-min fraction of input molecules bound, and (d) 15-min absolute captured count. The shaded 0.25–0.50-mm interval is a candidate conditional design window; the dotted line marks 0.40 mm. A short t_80_ for a weak-affinity antibody can correspond to a much lower equilibrium capture ceiling and therefore poor absolute output. Captured count is also affected by input volume, which grows with squared radius.

Wide-range kinetics at the representative 0.40-mm inner radius. Fixed 20-mm length, 1-nM analyte and 15-min incubation, k_on_ = 10^6^ M^−1^s^−1^. Density is the bound population per µm^2^ at the assay endpoint, whereas t_80_ is relative to each different equilibrium. Calculated values, not instrument LODs.The calculated equilibration times, 15-min surface capture densities, and recovery fractions across the affinity range are reported in Table 3.

**Table 3.** Affinity-dependent equilibration times, surface capture densities, and recovery at a fixed tubular reactor radius.

| $K_D$<br>(nM) | $k_{off}$<br>(s<sup>-1</sup>) | $t_{80}$<br>(min) | $\Gamma_{15}$<br>(molecules/$\mu\text{m}^2$) | 15-min recovery |
| --- | --- | --- | --- | --- |
| 0.01 | $10^{-5}$ | 16.17 | 93.75 | 77.8% |
| 0.1 | $10^{-4}$ | 15.96 | 93.05 | 77.3% |
| 1 | $10^{-3}$ | 14.14 | 86.54 | 71.9% |
| **10** | $10^{-2}$ | **6.70** | **49.99** | **41.5%** |
| 100 | $10^{-1}$ | 0.71 | 9.13 | 7.6% |

The central physical interpretation is not that weak affinity makes the reactor faster. At 0.40-mm radius the 10-nM antibody reaches 80% of its smaller equilibrium in only 6.70 min, compared with 15.96 min for the 0.1-nM antibody, while its 15-min surface density falls from 93.05 to 49.99 molecules/ µ m^2^. At 100 nM, the relative equilibration benchmark becomes shorter still (0.71 min) but the captured density collapses to 9.13 molecules/ µ m^2^. In this counterexample, optimizing t_80_ alone would systematically select the analytically inferior recognition input. The quantity Γ_15_/Γ_eq_ must not be confused with either binding-site occupancy Γ_15_/B or the final ART detection limit.

### Two controlled routes to higher affinity: an explicit 1-to-0.1-to-0.01-nM comparison

The wide-K_D_ scan alone does not isolate how affinity is improved. We therefore compared two tenfold steps, K_D_: 1 → 0.1 nM and 0.1 → 0.01 nM, using two independently controlled kinetic paths from an identical starting antibody (k_on_ = 10^6^ M^−1^s^−1^, k_off_ = 10^−3^ s^−1^, K_D_ = 1 nM). Path O holds k_on_ = 10^6^ fixed and reduces k_off_ by factors of ten; path F holds k_off_ = 10^−3^ fixed and increases k_on_ tenfold at each step. All six points are computed with the same mass-conserving reversible cylindrical-PDE solver used above, at *l* = 20 mm, r = 0.40 mm, C_0_ = 1 nM, t_e_ = 15 min and fixed accessible site density. The comparison changes only the nominated molecular rate constant, not geometry, reporter or detector parameters.

The equilibrium Γ_eq_ (K_D_) is identical on the two paths for a given K_D_ because the equilibrium occupancy depends on their ratio, not on the individual rates. Finite-time trajectories and t_80_ nevertheless differ. As summarized in Table 4, density gains are calculated as 100 [Γ_15_ (K_D_/10)/Γ_15_ (K_D_) − 1] within each fixed-rate path, not against a different baseline. The corresponding differences in finite-time capture density and equilibration kinetics between the two affinity-improvement pathways are illustrated in Figure 12.

**Table 4.**
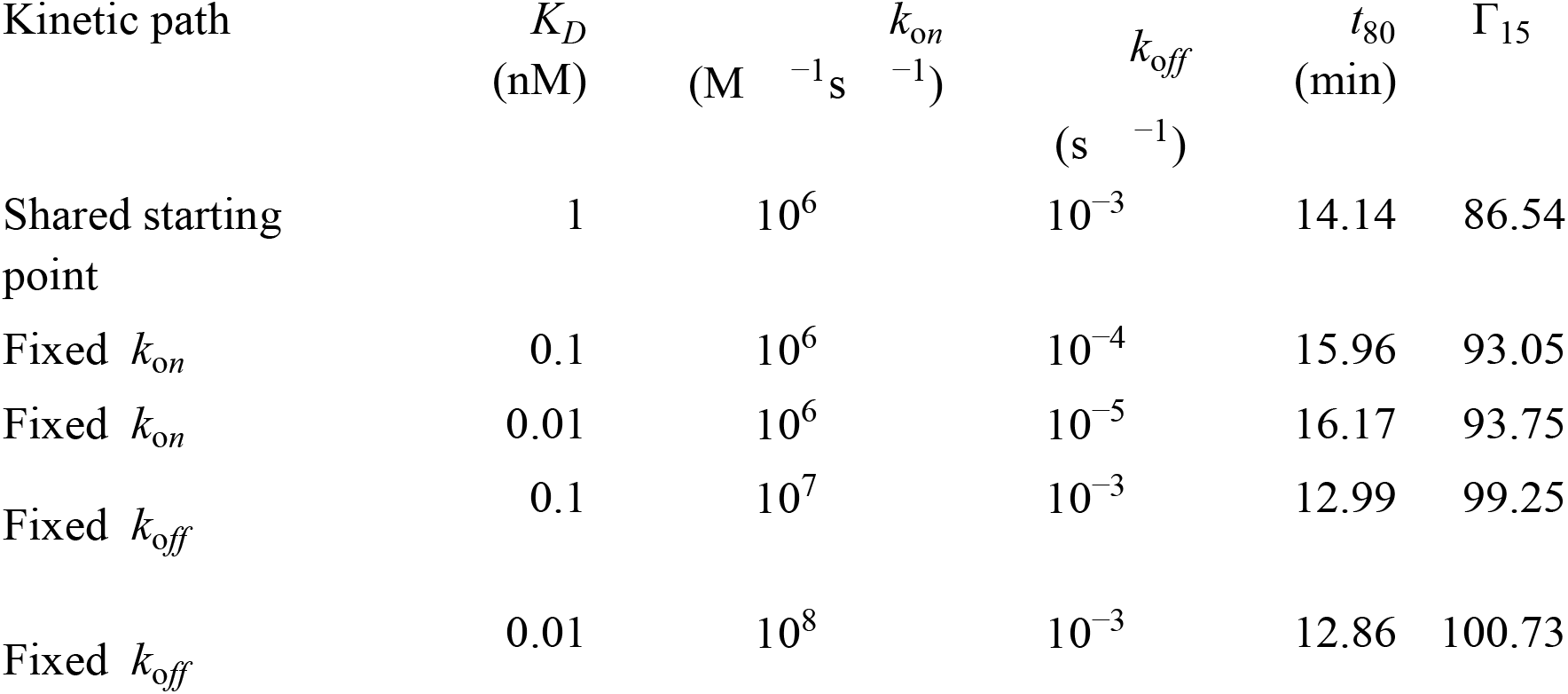
Comparison of two kinetic pathways for successive affinity improvements in a fixed tubular reactor.

| Kinetic path | $K_D$<br>(nM) | $k_{on}$<br>($\text{M}^{-1}\text{s}^{-1}$) | $k_{off}$<br>($\text{s}^{-1}$) | $t_{80}$<br>(min) | $\Gamma_{15}$ |
| --- | --- | --- | --- | --- | --- |
| Shared starting point | 1 | $10^6$ | $10^{-3}$ | 14.14 | 86.54 |
| Fixed $k_{on}$ | 0.1 | $10^6$ | $10^{-4}$ | 15.96 | 93.05 |
| Fixed $k_{on}$ | 0.01 | $10^6$ | $10^{-5}$ | 16.17 | 93.75 |
| Fixed $k_{off}$ | 0.1 | $10^7$ | $10^{-3}$ | 12.99 | 99.25 |
| Fixed $k_{off}$ | 0.01 | $10^8$ | $10^{-3}$ | 12.86 | 100.73 |

**Figure 12.**
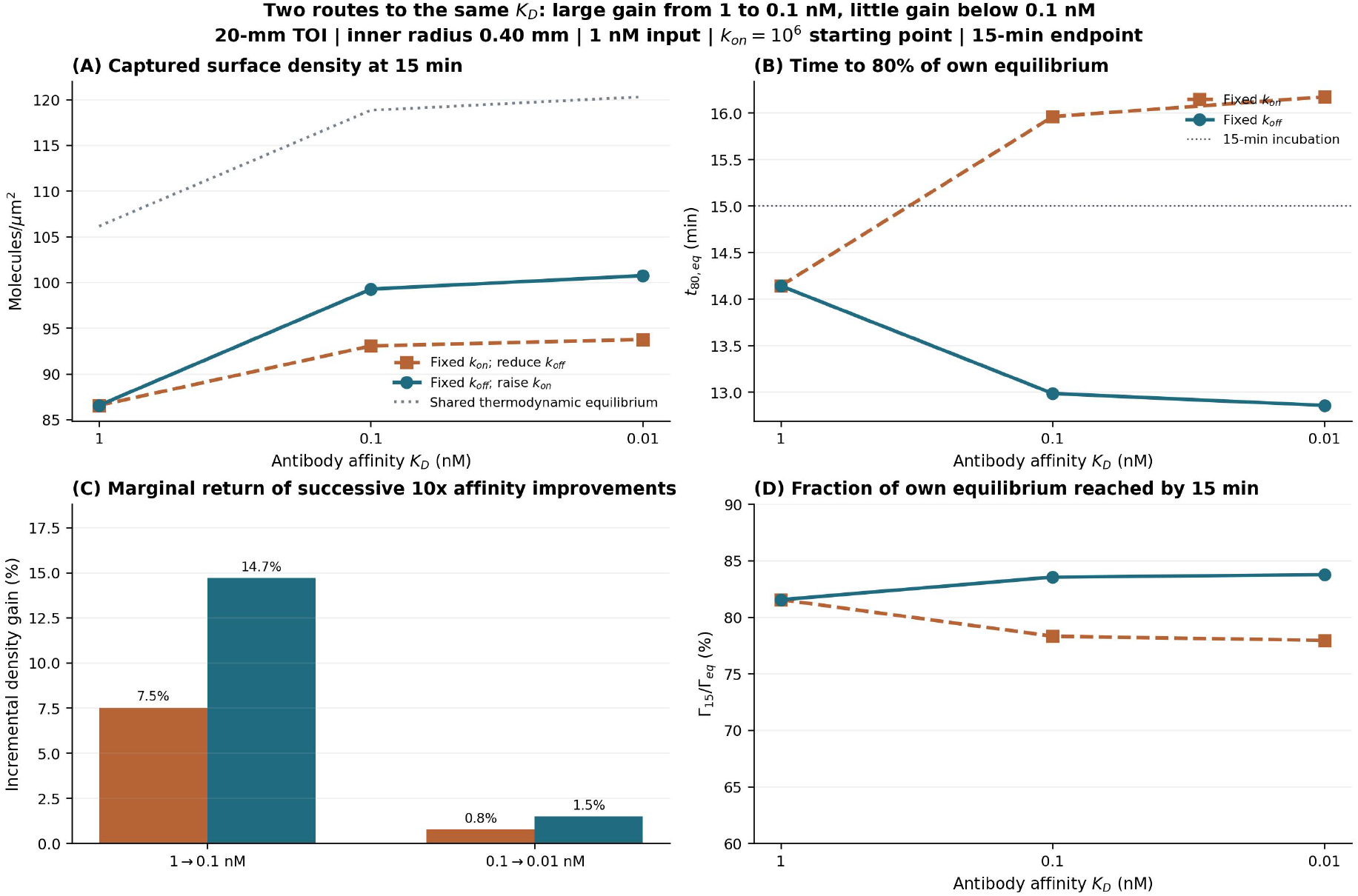
How affinity is improved matters more than K_D_ alone. (a) Fixed 15-min capture density for K_D_ = 1,0.1,0.01 nM under two distinct controlled perturbations, together with their common equilibrium surface density. (b) t_80_, the time to reach 80% of each condition’s own equilibrium density: lowering k_off_ at fixed k_on_ can prolong this relative time by increasing the equilibrium target, whereas raising k_on_ at fixed k_off_ shortens it. (c) The incremental density gain is +7.5% versus +14.7% for 1-to-0.1 nM, but only +0.75% versus +1.49% for 0.1-to-0.01 nM. (d) Fraction of equilibrium attained by 15 min. Fixed 20-mm length, r = 0.40 mm, C_0_ = 1 nM; theoretical AR populations, not measured sensor LOD.

Two successive affinity improvements at a fixed 0.40-mm TOI design. Source is a direct selection from the wide-rate sweeps, not a newly fitted model. Both paths share the starting 1-nM antibody and, at each K_D_, the same finite-bath equilibrium; they have different approach kinetics. Γ_15_ is captured molecules per µm^2^ after 15 min.

Crucially, the two paths give a different interpretation of the “good enough antibody” threshold. Improving 1 nM to 0.1 nM still increases 15-min density by 7.5% (lowering k_off_) or 14.7% (raising k_on_), while going on to 0.01 nM adds only 0.75% or 1.49%, respectively. More affinity is not a universal route to faster equilibration: t_80_ increases from 14.14 to 15.96 to 16.17 min on the fixed-k_on_ route, whereas it falls to 12.99 and then 12.86 min on the fixed-k_off_ route. These observations favor sufficient and sufficiently fast recognition over blindly minimizing K_D_, but do not imply that k_off_ is unimportant for longer washing, cross-reactivity, or matrix-dependent selectivity.

When does A or R dominate an AR-only detection threshold?

A/R dominance is meaningful only after specifying the output metric and the intervention constraints. In particular, a favorable tube-design effect on an AR threshold does not prove a larger effect on complete-instrument LOD. Define C_AR_ as the input concentration at which a stipulated number of retained target molecules (N_thr_ = 10^5^ here) is reached under the stated capture and idealized wash conditions; this is an AR-only concentration threshold, not the final T_m_/T_s_-constrained assay LOD. In the low-occupancy TOI simulation, compare equal fractional increases of k_on_ and equal fractional decreases in radius, while holding sample volume, dissociation rate and other assay inputs constant:

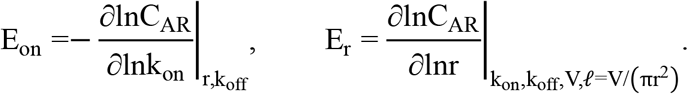

An A-dominant intervention is precisely E_on_ > E_r_: improving association kinetics lowers the threshold faster than the chosen reactor redesign. The opposite inequality gives an R-dominant intervention; the equality is a crossover, not a thermodynamic phase boundary. The full 5,304-point modeled r × k_on_ grid is plotted in Fig. 13. At the stated TOI reference E_on_ ≃ 0.13 and E_r_ ≃ 0.59, so the reactor redesign has about 4.6-fold larger leverage than the same fractional improvement in k_on_. At slower association rates (k_on_ ~ 10^4^ M^−1^s^−1^ near r = 0.40 mm), the association elasticity increases to about 0.95 while the radius elasticity is about 0.98: both controls matter, without a strong A-only advantage. In this particular fixed-volume comparison the strict E_on_ > E_r_ region is narrow, rather than an artificially large blue A-dominant territory.

**Figure 13.**
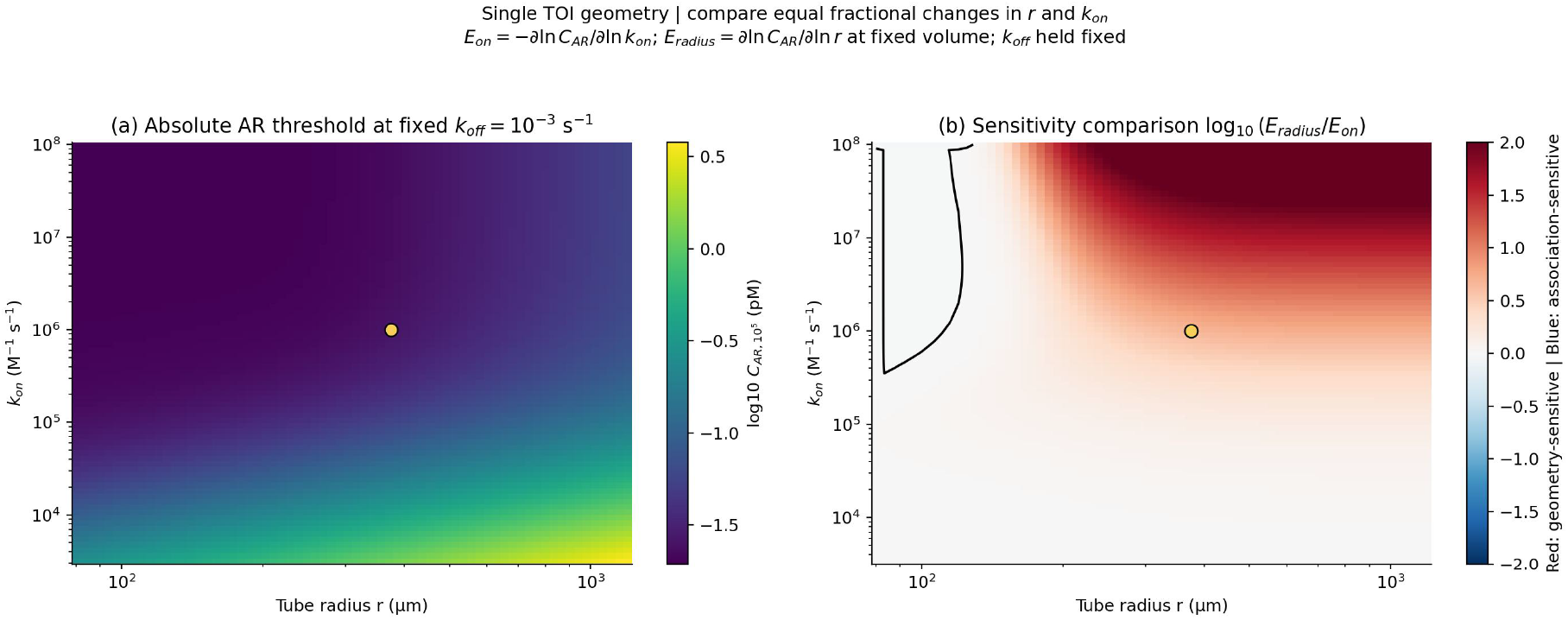
A/R control regimes can be defined by the leverage of two explicit design interventions. Left: the modeled input concentration needed to retain a stipulated 100,000 target molecules after a 15-min incubation and 30-s ideal-sink wash (AR-only threshold, not instrument-level LOD), versus tube radius and k_on_, with k_off_ = 10^−3^ s^−1^. Right: log_10_ (E_r_/E_on_) for matched fractional interventions, where E_on_ =− ∂lnC_AR_/∂lnk_on_ and E_r_ = ∂lnC_AR_/∂lnr along a fixed-volume radius/length redesign. Red signifies greater modeled leverage from reactor resizing, blue greater leverage from increasing k_on_, and the black contour is the crossover. At the TOI reference (gold marker; r ≈ 0.375 mm, k_on_ ≈ 10^6^ M^−1^ s^−1^), E_on_ ≈ 0.13, E_r_ ≈ 0.59 : reactor resizing has ~ 4.6 × the local leverage for this criterion. The blue recognition-dominant domain is narrow in this particular comparison. Changing radius also changes wall area and transport; the map does not isolate diffusion or directly quantify final ART LOD.

Figure 13 presents these elasticities as a conditional intervention map, not an instrumental LOD phase diagram

This control map asks a different question from the preceding fixed-20-mm design study. Here shrinking radius changes tube length *l* and binding surface area at fixed sample inventory, whereas the fixed-length study changes sample inventory with radius. Consequently the geometric leverage is not a measurement of “pure diffusion resistance,” and moving the A/R crossover will depend on which reactor variable is actually under design control. A downstream T_m_/T_s_ model may move the final LOD leverage boundary even when the AR threshold map is unchanged. The map therefore supplies the physically controlled A-versus-R evidence requested by the architecture, while avoiding a premature instrument-LOD claim.

For any two routes producing the same K_D_, an identical Γ_eq_ provides a strong internal control: differences in Γ_15_ and t_80_ are finite-time effects rather than a change in the thermodynamic endpoint. The full radius- and concentration-dependent affinity-gain comparison is retained in Supplementary Fig. S25. The highest association-rate condition (10^8^ M^−1^ s^−1^) is an illustrative kinetic boundary, not a claim that a specific available antibody possesses this rate. The finding concerns on-target AR capture. It cannot by itself establish a lower analytical LOD, because background AR populations, reporter loading, molecular transduction and photoelectric observation have not been changed or calibrated in this comparison.

Numerical audit: each finite-volume step conserves input-plus-surface molecules; across the 600-point wide-affinity sweep the maximum relative mass residual was 3.2 × 10^−12^. A further 360 fixed-k_off_ calculations were performed over nine on-rates, ten tube radii and four concentrations; their reported coarse-to-fine numerical checks remained below 0.02% in the selected conditions. A five-point coarse/fine grid/time-step stress check included the 10- and 100-nM conditions and found relative differences below 0.03% in t_80_ and 15-min density. These numerical tests establish discretization consistency under the specified homogeneous, quiescent reactor assumptions, not validation of kinetic rates, surface coatings, assay SNR or clinical analytical limits.

To express the broad useful range without forcing a spurious maximum of the wall density, we retain two transparent alternatives. First, define a conditional balanced utility 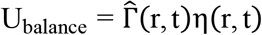, where 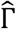 is explicitly normalized across the stated comparison range. This chosen equal-product weighting has a conditional interior peak, but the chosen weights are not a kinetic law or a calibrated analytical objective. Second, choose externally defensible performance constraints (minimum density, target capture fraction, finite assay time, specimen volume or maximum accepted t_80_) and report the feasible radius interval. The former makes one tradeoff visible; the latter more accurately describes an application-specific design decision. The existing FPT-surrogate peak remains an interesting approximate design-scale heuristic, not an exact prediction of the spatially resolved wall-density curve.

For the high- and intermediate-affinity reference inputs, the present results support treating inner radii of about 0.25–0.50 mm as a candidate high-utility design range, with 0.375–0.40 mm valuable practical reference points near a 15-min kinetic operating benchmark under the current k_on_ values. The expanded K_D_ = 10 and 100 nM results demonstrate that a comparatively short t_80_ can coincide with a markedly smaller captured density; neither the design window nor a ranking based solely on t_80_ is platform-independent. They do not establish that exactly 0.40 mm optimizes a particular optical instrument. Such a claim would additionally require spatial reporter generation, collection geometry, local background variance, and T_m_/T_s_ readout calibration.

At very low input concentration, deterministic mean density is insufficient: N_in_ is approximately Poisson distributed with mean c_0_πr^2^*l*, and 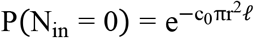. For a 20-mm tube at r = 100 µm, the expected input count is approximately 378 molecules at 1 fM but only 3.78 at 0.01 fM (10 aM); further reducing radius therefore risks entirely empty specimens. Shrinking a microreactor can improve capture efficiency yet impair sample presence probability when total volume becomes too small. This trade-off must be propagated to T_m_ and T_s_ before drawing conclusions about realistic detection limits.

Mathematical quality control and limits. The Bessel/Robin eigenfunction checks are independent of the finite-volume algorithm and confirm the same nonzero large-radius limit. Across 2720 reversible-PDE calculations, relative closed-system molecular mass-balance residuals did not exceed 2.9 × 10^−12^. A coarse-to-fine refinement changed 900-s results by less than 0.06% at representative radii, while the 30-s coarse-to-fine difference reached roughly 1%, establishing a more conservative precision bound at very early time. The analytic Robin comparison omits dissociation; the reversible PDE at 10 fM and K_D_ = 0.1 nM differed from irreversible Robin predictions by less than 1% after 900 s at the specified reference radii, consistent with the modeled off-rate. Neither these numerical checks nor the exact idealized diffusion result validates optical signal collection, nonuniform coating, actual antibody kinetics, mixing, wall reflections, sample heterogeneity or camera noise. Consequently, the appropriate statement is: Passive tubular confinement accelerates molecular recovery and enables low-volume, short-time operation; it does not universally maximize local captured surface density, and its actual LOD advantage requires full ART observability and noise analysis.

### Molecular transduction and observation: from captured complexes to reportable signal

Physical interface: AR delivers reporter-bearing molecular complexes. Tm converts their activity into a chemical product or time-dependent photon emission; Ts constrains which of those physical outputs becomes the reported measurement. This section introduces the minimum effective equations needed for the full chain. ART fixes the order AR → Tm → Ts, not one chemistry-specific formula. The following enzyme law is an explicit, compact candidate for a substrate-limited Tm, rather than an assertion that all reporters obey Michaelis–Menten kinetics. It is derived below; its numerical calibration and cross-condition validity remain to be tested. The effective Ts model intentionally uses an externally specified detection threshold instead of a detailed noise budget.

### T_m_: a four-parameter, substrate-conserving conversion function

Let N be the effective number of enzyme-bearing complexes delivered by AR after label binding and washing. Convert this inventory to active-enzyme concentration as E = αN, where α includes the number of active enzyme equivalents per reporter and the reaction volume (e.g., mol L^−1^ per complex). For a chosen effective limiting substrate S (t) with known starting concentration S_0_, approximate the reaction progress by

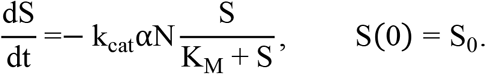

The assumptions are intentionally explicit: a well-mixed reaction with constant active enzyme, one effective limiting substrate, an approximately fixed turnover yield and no dominant enzyme inactivation or product inhibition over the chosen development interval. HRP–TMB–peroxide and HRP–luminol–peroxide are multi-reactant chemistries; S_0_ and K_M_ here must therefore be understood as effective quantities unless the controlling reagent is independently identified. The equation is not an exact mechanism for all two-substrate regimes. The integrated substrate-depletion approach follows standard enzyme-progress reasoning.[13]

Define a dimensionless consumed-substrate fraction f (N, t) = 1 − S (t) /S_0_, constrained to 0 ≤ f < 1. Integration of Eq. [eq:Tm_MM] gives

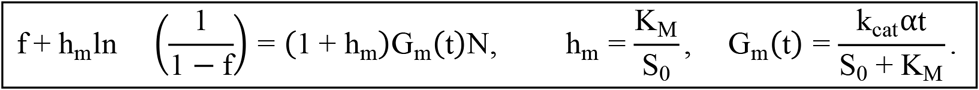

This is a one-variable implicit algebraic equation, not another transport PDE. At a specified reaction time, the accumulated reporter response is the four-parameter map

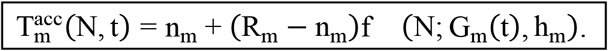

Here n_m_ is the chemical baseline (not a detector floor), R_m_ is the molecular product-capacity roof set by available limiting reagent and conversion yield, G_m_ is the initial normalized response per effective enzyme-bearing complex, and h_m_ = K_M_/S_0_ fixes the shape of substrate depletion. These quantities are meaningful only for a specified chemistry, reaction volume, time and observable. G_m_ is dimensioned so G_m_N is dimensionless, and varying t changes G_m_; the four parameters are not freely re-fitted for each experimental time point.

Two mathematical checks make the physical interpretation precise:

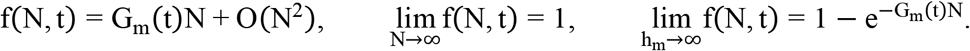

For finite h_m_ > 0, the response approaches its roof smoothly, not through a hard corner. The incremental response falls continuously; h_m_ controls the curvature in a chemically interpretable way. A small h_m_ = K_M_/S_0_ means an initially nearly zero-order, less-curved depletion trajectory with a comparatively concentrated final transition; large h_m_ approaches gradual first-order exponential depletion. In the low-input expansion, 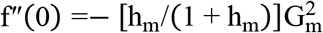, while f′ 0 = G_m_. Thus the meaning of h_m_ does not depend on inventing an arbitrary Hill exponent.

The constraint across conditions is as important as the fit to one curve: at fixed reagent chemistry, h_m_ = K_M_/S_0_ changes predictably with the starting limiting-substrate concentration, G_m_ (t) ∝ t/ (S_0_ + K_M_), and the chemical product roof above baseline scales with the available convertible substrate in the idealized limit. The response depends on enzyme–time dose Nt when S_0_, K_M_ and catalytic activity are held fixed. These are predictions to test, not new experimental observations. The predicted substrate-conversion dynamics and their implications for accumulating product and time-dependent photon emission are illustrated in Figure 14.

**Figure 14.**
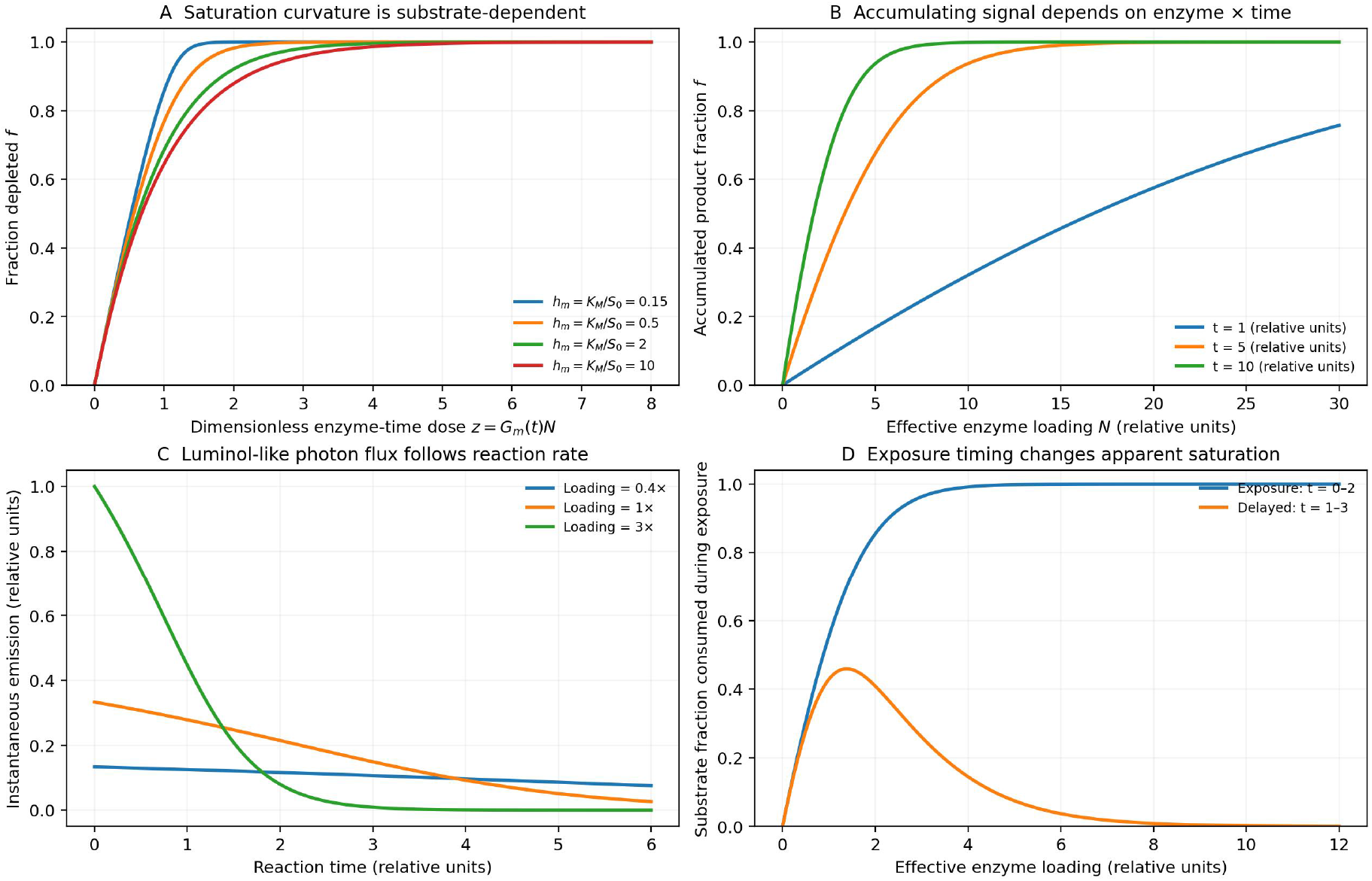
Substrate capacity links saturating color and transient light emission. (A) The dimensionless consumed-substrate fraction from Eq. [eq:Tm_progress] approaches a reagent-defined capacity continuously for different h_m_ = K_M_/S_0_. (B) Changing reaction time translates the accumulating response through the enzyme–time product. (C) The same reaction progress gives an instantaneous, decaying photon-generation rate for a hypothetical luminol-like readout. (D) A delayed exposure can yield a non-monotonic photon count even when exposure starting at t = 0 is monotonic. Curves are dimensionless simulations with arbitrary normalized enzyme and time scales, not measured TMB or luminol data.

### Two temporal outputs of enzyme conversion; direct fluorescence as a separate limit

A conserved conversion-progress variable supports two distinct experimental output types without separate saturating response laws. For accumulating product (stopped TMB or a stable enzyme-generated fluorophore), the product concentration at development time t_d_ is proportional to S_0_f N, t_d_ under constant stoichiometric yield. Following acid stop, the appropriate TMB chromophore concentration enters a wavelength-specific optical measurement; the optical readout roof must not be confused with the chemical product roof.

For real-time chemiluminescence, the molecular product is instead a photon-emission rate. Under a fixed effective chemiluminescence photon yield ϕ_CL_ per consumed limiting-substrate molecule, reaction volume V (L), and Avogadro constant N_A_,

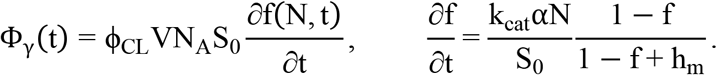

The photon total emitted during a specified acquisition interval [t_a_, t_b_] is

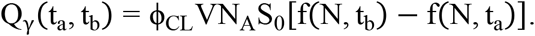

The conversion rate Φ_γ_ t is the T_m_ output; the time integration into detected photoelectrons belongs to T_s_, not to chemical accumulation in the sample. Equation [eq:Tm_CLwindow] is the corresponding chemically emitted photon budget over a chosen interval, not a second, independently fitted cumulative reporter law. If acquisition begins when substrate is added (t_a_ = 0), accumulated emitted photons increase toward a finite chemical capacity with increasing N. If it begins after a delay (t_a_ > 0), a large enzyme load may consume much of the substrate before the window starts; collected photons can then decline as N increases. A monotonic four-parameter endpoint sigmoid is consequently inappropriate for every delayed-read luminol experiment. HRP–luminol emission may additionally show induction, enhancer effects and multiple kinetic regimes; published time-resolved models and data confirm that its full kinetics can be more complicated than this effective depletion law.[14,15]

Directly conjugated fluorophores do not require a consumable enzyme substrate. Within their unsaturated excitation range an appropriate, even simpler T_m_ is Φ_F_ t ≃ β_F_NI_exc_ (t), where β_F_ groups fluorophore loading, absorption and quantum yield. Photobleaching or excitation saturation can be introduced only when material for a particular experiment. The four-parameter enzyme law is therefore a replaceable family member, not a compulsory law for every label.

### T_s_: an operational floor–slope–roof observation rule

For most conventional immunoassay readers—photodiodes, PMTs operating within their linear range, and CCD/CMOS imaging—a proportional observation law is sufficient over the useful window. Define M_s_ = *O*_s_ [T_m_] as the physical reporter quantity selected by the assay readout: stopped optical absorbance or emitted photons over a defined exposure, with required wavelength/time integration treated as known parts of the observation procedure. Let g_s_ convert that quantity to the instrument’s reporting units. The minimal final reporting model is

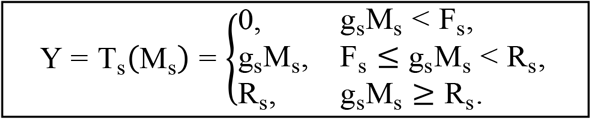

F_s_ is a specified effective detection floor and may deliberately be set conservatively from blank measurements or operating practice; g_s_ is the linear observation slope; R_s_ is the detector/readout roof. The reported zero beneath F_s_ means not distinguishable for this assay protocol, not that zero photons or zero bound molecules exist. The model does not require deriving shot noise, read noise, dark current, false-positive distributions or specialized avalanche-detector kinetics; such descriptions may replace this T_s_ implementation if a particular experiment requires them.

For emitting reporters, a standard physically interpretable slope is based on collection, optical throughput, quantum efficiency, exposure timing and electronic conversion. Before the piecewise reporting rule, the expected number of collected electrons can be represented as

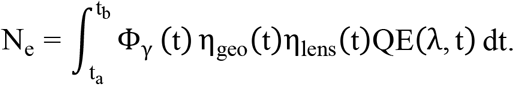

For colorimetry, the input state is a chromophore concentration produced and stopped by T_m_. In the working Beer–Lambert regime the wavelength-specific readout obeys

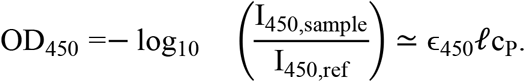

A validated upper OD450 reporting range, a chemical substrate roof, pixel ADC capacity and binary digital saturation remain distinct physical limits. In particular, a nominal upper validated OD of 2.0 corresponds to roughly 1% transmission, but is neither a universal TMB chemical ceiling nor a statement about sensor full-well capacity.

### Background, digital readout and pragmatic verification

True and false AR reporter-bearing complexes are both passed to T_m_; increasing enzymatic gain does not retroactively distinguish them. Background may be represented through a specified effective chemical baseline n_m_ and a conservative downstream floor F_s_. Here we do not construct a separate noise variance decomposition: T_s_ records zero whenever the pre-threshold signal fails the adopted decision floor, and otherwise follows the linear slope until the roof. The inherited illustrative background curves below are useful context, not parameters derived for the new enzyme law. The operational observation floor and the reporting of subthreshold signals are illustrated in Figure 15.

**Figure 15.**
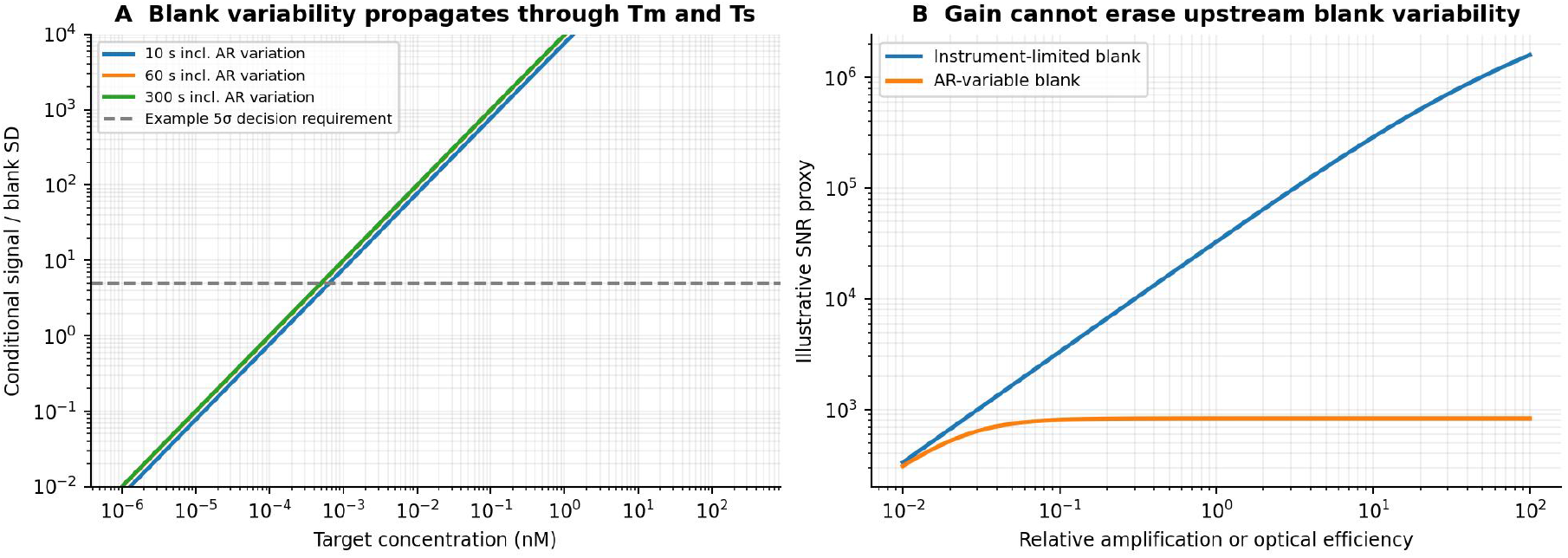
Molecularly generated signal can remain below the observational floor. An inherited illustrative floor diagram. The adopted T_s_ operator is intentionally an operational hard threshold, not a derived noise distribution: subthreshold generated signal is reported as zero.

For partitioned digital immunoassays, a separate ideal occupancy-statistic limit is p_+_ = 1 − exp [− (λ_T_ + λ_BG_)]. Binary positive fraction can approach unity while analog photon/electron measurement has considerable remaining physical headroom. These are different upper limits, not an argument for treating digital saturation as a molecular or detector roof. At p_+_ = 99.99%, ideal occupancy λ ≈ 9.21. In one inherited illustrative calibration, this corresponds to about 0.194 analog units against a 1000-unit detector roof, or approximately 3.7 decades of remaining analog headroom; the numerical difference is calibration-specific.

A lean prospective validation route, which remains to be completed, is to characterize enzyme activity and limiting-substrate conditions independently, then test whether the same S _0_, effective K _m_, yield, and enzyme loading reproduce TMB responses across different development times and substrate concentrations. For luminol, compare instantaneous light curves and two early/delayed acquisition windows under a controlled reagent protocol. For a direct fluorescent label, check the linear unsaturated branch. The strongest test is a prediction after changing time or reagent concentration without refitting all four parameters; agreement for one curve alone does not establish transferability. No such prospective experimental validation is claimed in this manuscript.

### End-to-end ART illustrations: legacy ELISA and tubular chemiluminescence scenarios

Closure test: Two assays may use the same intrinsic A parameters without sharing reactor physics, reporter time evolution, or optical observation. These inherited, conditional examples demonstrate the interfaces, not independent experimental validation of the newly proposed four-parameter T_m_ law. The two case studies are not the same AR followed by different reporters. Buffer comparisons and cross-reactivity above are conditional tests at a fixed plate; the end-to-end comparison below instead transfers a common baseline recognition parameter set into two independently calculated reactor environments. They share the illustrative antibody parameters k_on_ = 106 M^−1^s^−1^, k_off_ = 10^−3^ s^−1^ (K_D_ = 1 nM) and target concentration scale, but calculate AR (A, R_E_) and AR (A, R_T_) independently. Effective surface capture probability, reagent loading, wash and reporter chemistry may in real experiments modify apparent rates; their transferability across supports is a hypothesis requiring measurement. The cases explicitly state controlled simplifying assumptions rather than claiming the complete multilayer sandwich assay has been fitted.

### A common recognition input gives different reactor-specific AR states

Using the same reference reduced law, 1 nM input at 10 minutes predicts Γ_E_ = 91.80 (plate) and Γ_T_ = 75.38 (TOI) molecules/ µm^2^. At 15 minutes these become 113.31 and 87.20, respectively. With reactive areas of 32.0 and 47.12 mm^2^, respectively, the latter endpoint corresponds to X_E_ ≃ 3.63 × 10^9^ and X_T_ ≃ 4.11 × 109 bound molecules. The two absolute numbers cannot be equated to recovery fraction or a reporter count. For these legacy illustrative full-chain scenarios (which do not validate Eq. [eq:Tm_progress]), the detector-antibody reporter competency is approximated by an identical conditional probability p_det_ = 0.5 and a 30-s no-rebinding wash; in a predictive two-step assay that step must instead be solved as a second reactor-specific AR operation.

### Illustrative Case I: 96-well plate → HRP–TMB → stopped OD450

The plate captures target with its own AR_E_, followed by HRP-labeled detection antibody and wash. After adding TMB and peroxide, HRP catalysis forms colored product during one of three separate development periods (1, 5, 10 min). Acid stopping terminates the reactions and yields a yellow chromophore read at a specified λ = 450 nm. This is an endpoint OD450 protocol. An unstopped blue TMB reaction read at approximately 620–650 nm is a different experiment and is not substituted into this case [3,4].

With an effective enzyme-label count N_E_ C (target plus false-event labels), the source conditional example uses the finite chemical-product reduction

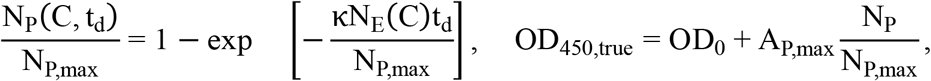

followed by the wavelength-specific T s observation in Eq. [eq:od]. This inherited exponential example is the h_m_ → ∞ mathematical shape limit of Eq. [eq:Tm_progress] at fixed normalized gain, after matching the old rate normalization. It is not evidence that the actual TMB reagent has infinite K_M_/S_0_. It has not been recalculated for finite h_m_ = K_M_/S_0_. The assumed A_P,max_ = 3.0 OD-equivalent, effective rate normalization (2.5× 10^9^ labels per five minutes), labeling probability and background are illustrative inputs, not experimentally identified k_cat_, actual substrate stock or molar extinction coefficients. The finite N_P,max_ and OD450 working threshold have deliberately separate meanings.The resulting concentration-response curves illustrate the trade-offs between development time, low-concentration detection, and high-concentration saturation in the TMB–OD450 workflow (Figure 16).

**Figure 16.**
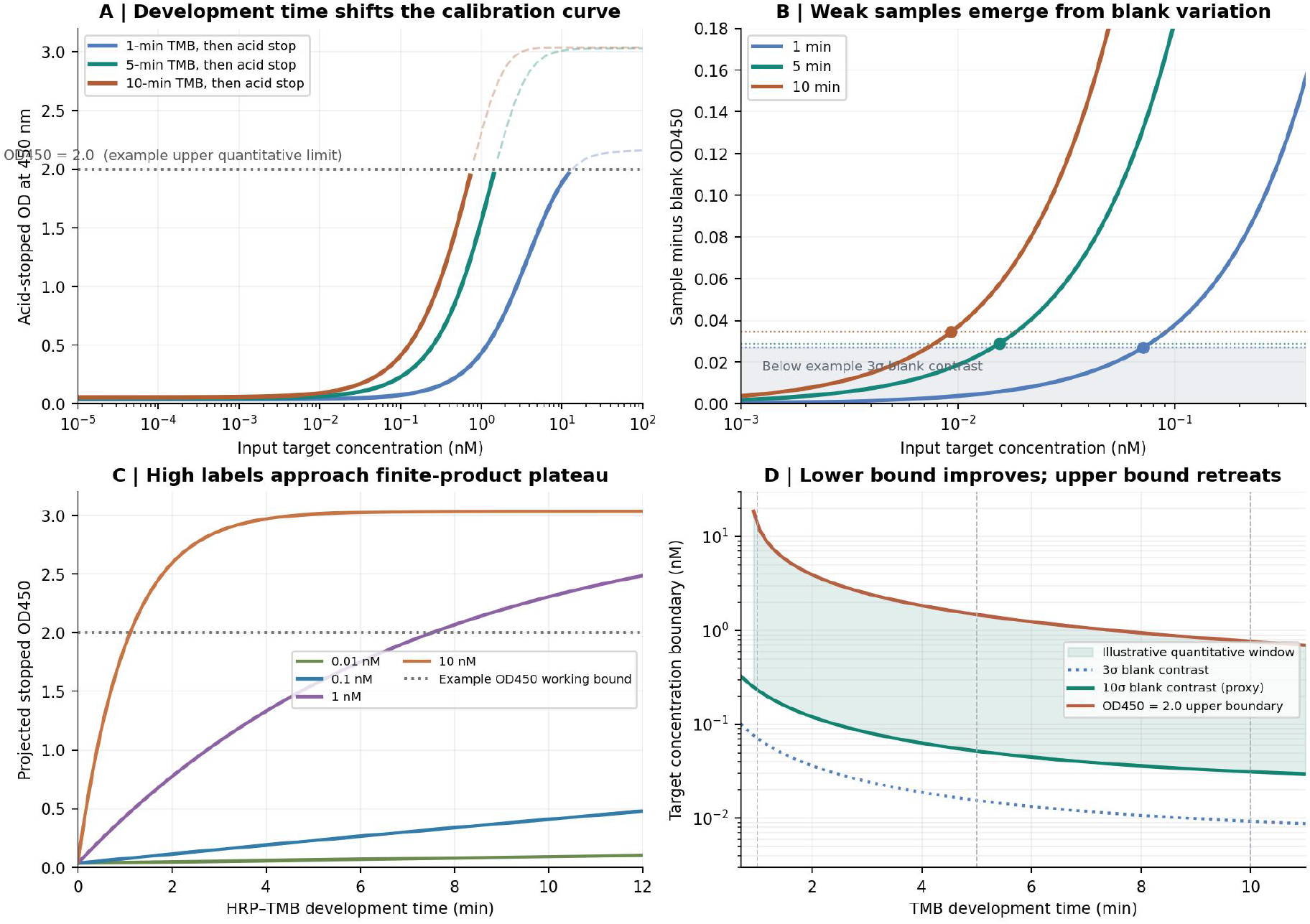
TMB incubation trades low-end access for high-end chemical/OD limits. The panels compare the full concentration response, an expanded low-end view, a high-label product plateau and the time-dependent reporting thresholds. Longer TMB development allows a lower analyte concentration to cross an inherited blank-based operational threshold but moves high concentrations toward finite-product saturation and beyond an illustrative OD450=2.0 validated reading boundary. Values exceeding the OD reporting limit are visually distinguished from hard clipping.

The inherited illustrative calibration, not a requirement of the deliberately simple main T_s_ law, uses an assumed AR-background enzyme population (8 × 10^6^ mean labels, 3 × 10^6^ SD), chemical/reagent contribution (OD SD 0.004) and reader contribution (OD SD 0.008). A first-order blank approximation is 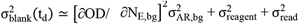. The calculated trade-off between blank response, concentration thresholds, and the upper OD _4 5 0_ reading limit across the three development times is summarized in Table 5.

**Table 5.** Predicted blank responses and concentration thresholds at different TMB development times.

| $T_m$<br>time | Blank<br>OD450 | $3\sigma$ threshold<br>(nM) | $10\sigma$ proxy<br>(nM) | OD450=2 crossing<br>(nM) |
| --- | --- | --- | --- | --- |
| 1 min | 0.037 | 0.07135 | 0.2339 | 13.52 |
| 5 min | 0.045 | 0.01554 | 0.05201 | 1.48 |
| 10 min | 0.054 | 0.009285 | 0.03127 | 0.7644 |

These legacy 3 σ and 10 σ thresholds are optional examples of how one could choose a conservative effective floor F_s_, not additional statistical machinery required by Eq. [eq:Ts_floorroof]. The concentration thresholds and OD upper crossing are outputs of a transparent hypothetical calibration, not measured ELISA LOD/LOQ and not recalculated from the new finite-h_m_ substrate law. A 10σ threshold is a working contrast proxy, not a regulatory or universally valid LOQ. All three develop times shift the blank as well as the target response. Apparent high-dose flattening may arise from chemical substrate depletion, loss of response slope near finite product capacity, or optical loss of validated OD range; it must not be described as one common saturation mechanism.

### Illustrative Case II: TOI capillary → HRP–luminol → side-view long-exposure camera

TOI uses its independently calculated AR_T_ and an HRP/luminol/peroxide chemiluminescent T m. At any time the emitted photon flux depends on active enzyme sites, kinetic turnover, finite reagent supply, reagent decay, and chemiluminescence quantum yield Φ_CL_. A simple exposure-integrated scenario model is

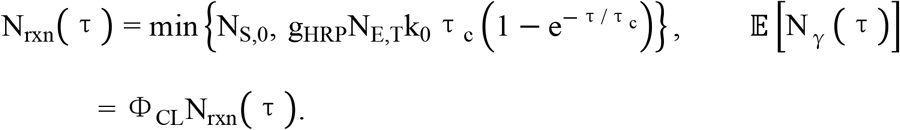

This inherited finite-pool, exponential-decay case is a phenomenological illustration and is not algebraically identical to the substrate-progress model in Eqs. [eq:Tm_progress] – [eq:Tm_CLwindow]; its photon curves have not been recalculated from the latter. The effective k_0_ = 1 reaction s^−1^ label^−1^ and τ_c_ = 90 s are illustrations only. PolyHRP alters g_HRP_ and the emitter production process within T m, while longer exposure changes T s. A calibrated study reported an optimized HRP – luminol – H _2_ O _2_ Φ_CL_ = 1.23 ± 0.20% under specified chemical conditions [5]; it is not a universal luminol quantum yield or a measurement of the present TOI reagent.

For the stated side-view optical scenario, approximate the emitting region initially as an isotropic point source centered 100 mm from a circular effective entrance pupil of diameter 70 mm. The geometric solid-angle collection factor is

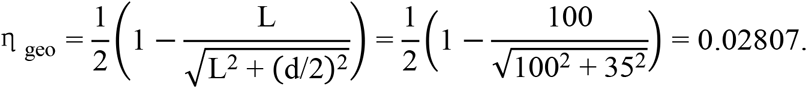

With η_lens_ = 0.80 and QE = 0.70 (both scenario inputs), the expected electron yield per luminol turnover is Φ_CL_ η_geo_ η_lens_ QE ≃ 1.93 × 10^−4^ in this scenario. A 70-mm external lens diameter is not automatically a 70-mm entrance pupil; tube-wall refraction, actual NA, emissive wall spatial distribution, shadowing and sensor spectral QE must be integrated or measured before assigning this collection fraction to hardware. The assumed geometric photon-collection efficiency is illustrated in Figure 17, while the predicted effects of exposure duration and effective HRP loading on the detected photoelectron yield are shown in Figure 18.

**Figure 17.**
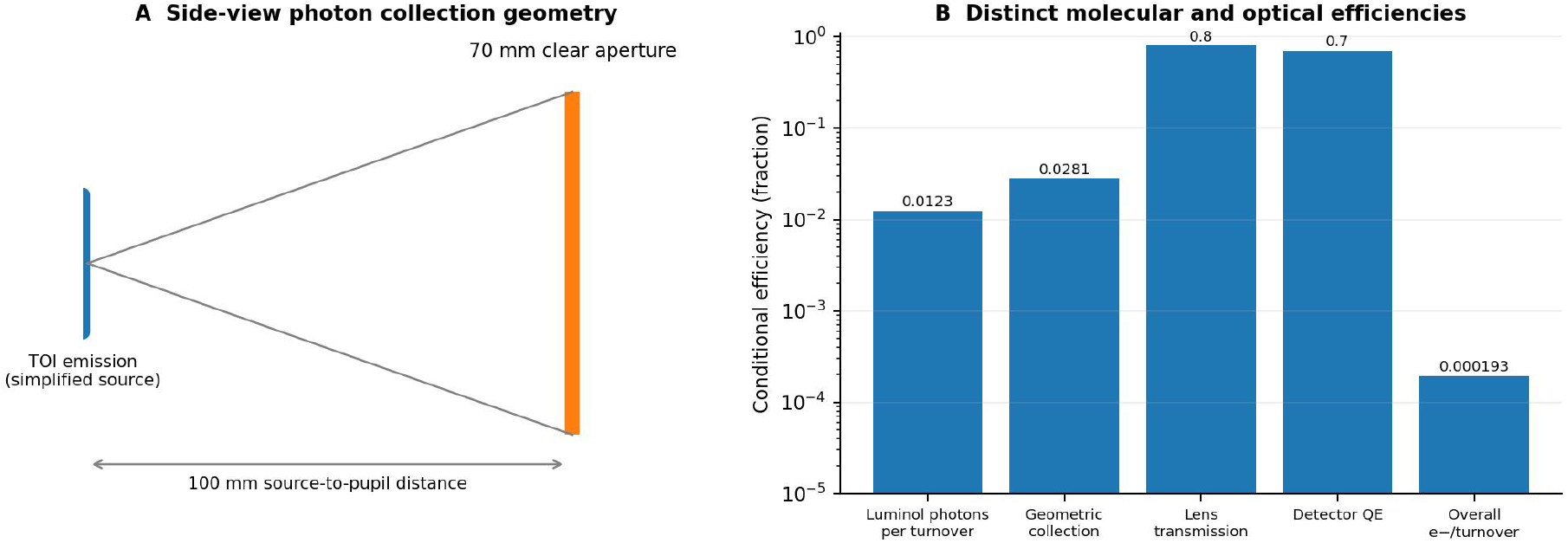
TOI CL is further constrained by geometric photon collection. 10-cm object-to-entrance-pupil distance, hypothetical 70-mm clear aperture, optional lens transmission and 70% sensor QE. The 2.81% geometric fraction is for a point isotropic source; real extended capillary emission and tube-wall optics require ray tracing or calibrated measurement.

**Figure 18.**
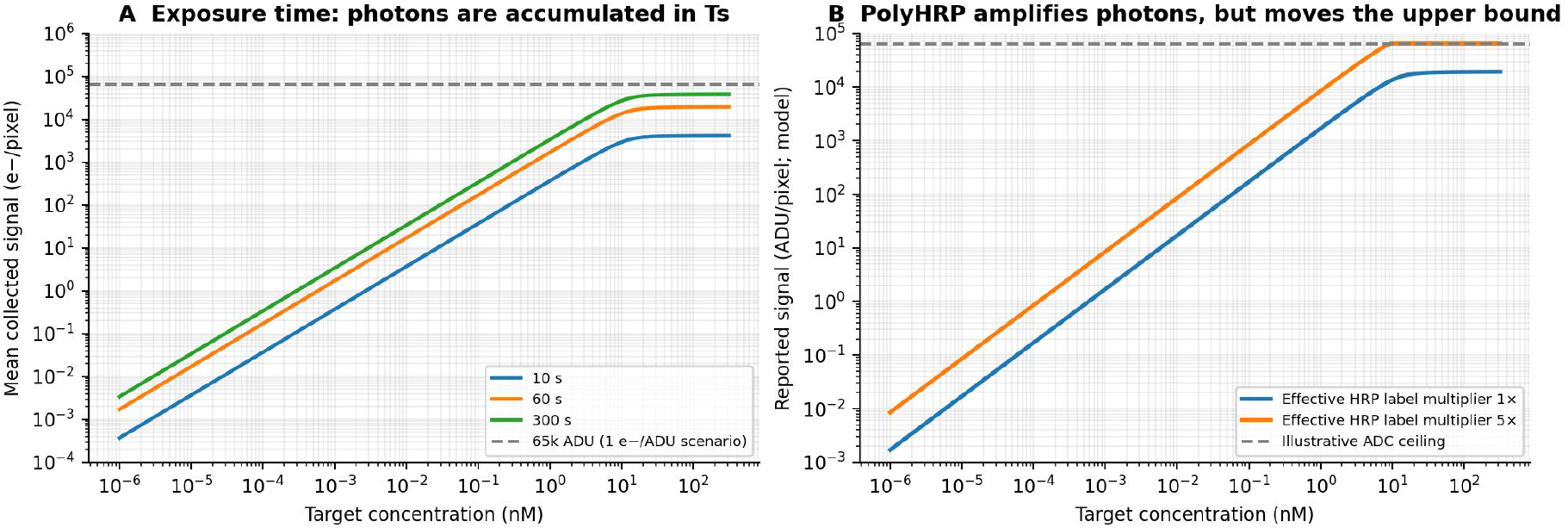
Longer exposure and reporter amplification compete with observation roof. Predicted photoelectrons for 10-, 60- and 300-s exposures, and a 1×/5× effective HRP loading contrast. The assumed 10,000-pixel uniform ROI and 1 electron/ADU are unmeasured display controls. A 65,000-ADU limit pertains to an individual pixel’s digital channel, not the integrated ROI photon inventory.

A camera display near 65,000 counts is compatible with a per-pixel 16-bit ADC limit (65,535 ADU) but does not establish full-well capacity, gain, spectral QE or the aggregated ROI headroom. Individual bright pixels can saturate before integrated ROI sums reach an arbitrary total. Long exposure increases signal, reagent and dark-background counts together; chemical photon generation occurs contemporaneously, not as an accumulating optical substance in the tube.

### Where the full chains gain and lose information

For ELISA, the dose-response axis passes through geometry-dependent occupancy, reporter antibody binding, finite HRP product accumulation, acid stop and fixed-wavelength OD450. The gain-time trade-off compresses high concentrations even as new low concentrations cross the blank threshold. For TOI, the dose response passes through a separate surface inventory, luminol photon emission, roughly percent-scale chemical quantum yield, percent-scale geometric collection, QE, exposure and pixel electronics. Both chains may amplify off-target enzyme labels. Both can lose low-end observability below a conservatively specified instrument floor. Neither can be inferred from the AR density curve alone. The existing numerical figures in these two cases document their original conditional assumptions; a new, prospectively calibrated application of Eq. [eq:Tm_progress] is the next step, not something already established by these plots.

## Discussion: what ART unifies, what remains conditional, and how to validate it

### The “perfect antibody” idealization

Under a simplifying optical-sensor view, the recognition element binds only the intended molecule, produces no off-target or nonspecific signal and never dissociates (k_off_ → 0 for target, false-event rates → 0). Then observed improvements are attributed chiefly to reactor or detector architecture. That is a legitimate local control model but removes chemically generated blanks by assumption. It does not make transport instantaneous, increase input analyte inventory, or turn an instrumental gain change into improved molecular selectivity. ART tests the difference between that counterfactual and finite cross-reactive capture.

### The “perfect confinement” idealization

Reducing diffusion distance can accelerate wall arrival (τ_D_ ~ L^2^/D) and enhance recovery, but arrival is not retained reporter-competent capture. Finite site inventory, off-target kinetics, reversible dissociation and sample volume remain. The same small capillary can increase recovery fraction and reduce absolute captured molecules under a fixed-length redesign. Consequently high S/V cannot serve as a universal substitute for the joint AR calculation.

### The “perfect transduction/observation” idealization

In an ideal readout, every physically informative event is measured with negligible transduction loss and detector noise. For a real-time single-molecule instrument, time-resolved event trajectories may provide additional kinetic information, and ART does not deny that discrimination capability. But in a washed endpoint assay, target, cross-reactant and nonspecific events may yield indistinguishable enzyme labels. Perfectly counting a false event is not a chemically correct assignment. Conversely, with nonideal T s, even real upstream light may never exceed the floor. These idealizations are conceptual limiting cases, not official names for the diverse fields of antibody engineering, microfluidics or single-molecule sensing.

### Why a single reactor yields a more interpretable design rule

The direct 1-to-0.1-to-0.01-nM calculation illustrates that a common equilibrium affinity does not specify the assay trajectory: at fixed off-rate, a faster on-rate improves the finite-time capture response more than an equal affinity improvement achieved solely by reducing off-rate in the reference TOI, and both routes show rapidly diminishing capture gains below 0.1 nM. This candidate threshold is conditional on target concentration, wash, time and geometry, and must not be promoted to an instrument-level LOD specification. Cross-platform rankings become misleading if surface density, total captured inventory and final analytical LOD are treated as interchangeable. The independent exact Bessel/Robin audit identifies a false large-radius density decay introduced by the MFPT single-exponential survival closure; full reversible-PDE short-time and low-concentration sweeps instead predict a monotonic-to-plateau local density under the specified passive tube assumptions, while narrow tubes improve finite-time fractional capture and sample economy. The density threshold remains explicitly separated from downstream chemistry and optical background. It predicts that shrinking the capillary radius at fixed sample volume may improve target collection more than further increasing k_on_ near the selected TOI operating point. But this comparison includes a longer tube and greater wall area; it is not a claim that pure diffusion resistance alone fixes TOI performance. A fixed-radius experiment varying mixing or diffusivity would be needed to isolate mass-transfer control directly.

The low-K_D_ plateau likewise depends on how affinity is improved. At fixed k_on_, reducing k_off_ from the K_D_ = 0.1 nM to 0.01 nM range changes the theoretical low-occupancy concentration threshold by only about 1%. Better k_on_ along a fixed-k_off_ path is a distinct intervention and can improve short-time capture in other parts of the map. A “good enough” antibody is therefore a conditional joint A × R × t concept, not a universal affinity threshold. Chemical cross-reactivity and blank noise may also improve through molecular engineering even when target-only binding yield has saturated.

For the previously considered sparse-bead setup, the high bound surface density was an unreliable surrogate for whole-assay performance; the stipulated target-count audit actually placed its theoretical concentration threshold above TOI’s. Plate and TOI target-count thresholds were superficially similar in part because their sample volumes differed more than tenfold. These limitations motivate geometry- and measurement-matched experiments before any direct ranking of final analytical LOD. Even for the TOI-only predictions, independent HRP loading, T m reporter yield, T s photon statistics, real blank distributions and wash-rebinding calibration remain prerequisites for a validated detection limit.

### The central intellectual contribution: a reusable physical coupling architecture

ART’s defining proposal is the structural identity Y = T_s_ [T_m_ [AR (X)]], understood as a composition of physically parameterized, potentially stochastic operators. Its originality cannot rest on the notation alone, because function composition, mass-action kinetics, diffusion, enzymatic amplification and detector statistics are established tools. The substantive claim is a physically reasoned partition of the analytical chain: A and R form a single reaction–transport state generator rather than sequential capture gains; T_m_ maps those molecular populations into chemistry-specific reporter states; and T_s_ maps reporter states into measured data. This partition establishes meaningful substitution points while preserving upstream backgrounds and downstream-added noise. It specifies what a complete model must exchange across interfaces and what cannot legitimately be decoupled.

The AR developments in this paper serve as a particularly detailed demonstration of that claim. Same-K_D_ families, casein/PBST perturbation hypotheses, spatial PDE audits, wash/rebinding, tube-dimension tradeoffs and t_80_ show why a model of recognition alone or geometry alone cannot define finite-time assay capture. For the specified TOI reference, increasing affinity from 1 to 0.1 nM can materially improve capture, while further improvement to 0.01 nM often gives much smaller marginal gains. That result is conditional on the input concentration, the on-rate/off-rate improvement path and the reactor. It is an AR demonstration of diminishing returns, not a universal antibody specification or a stand-alone prediction of analytical LOD.

The current AR formulation has reached a provisionally usable theoretical state, subject to the declared limits of the PDE, simplified transport clock, independently checked approximations and unmeasured experimental parameters. The T_m_ and T_s_ sections remain replaceable, chemistry-dependent working implementations. The enzyme branch now derives its four effective parameters from limiting-substrate capacity, enzyme-time gain and h_m_ = K_M_/S_0_, while accumulated product and real-time photon emission are alternate outputs of the same reaction-progress law. Ordinary observation is reduced to a practical floor–slope–roof rule, with the floor selectable conservatively. The assumptions behind the effective substrate, reporter yield, labeling stoichiometry and optical calibration still require independent tests; full detector-noise or specialized counting models are not required to make the ART composition usable. Such revisions are not exceptions to ART: model interchangeability under explicit physical interfaces is the point of its compositional design.

A stringent next test is prospective full-chain prediction. Characterize recognition and cross-reactivity, reactor geometry and accessible surface sites, limiting-substrate chemistry, effective molecular conversion, and observation slope, floor and roof independently; then predict an assay’s reported signal as time, substrate stock and reactor conditions change, with no assay-specific readjustment of hidden gains. This separates the theoretical contribution—the coupling map and the admissible module interfaces—from the much stronger empirical claim of universal quantitative predictiveness, which remains to be established.

## Conclusion

ART is proposed as a compositional physical theory of endpoint solid-phase immunosensing, organized by the coupling architecture

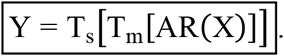

Its principal intellectual contribution is to define the physical division of labor and admissible interfaces along the entire analytical chain. Molecular recognition and reactor physics are inseparable when predicting the finite-time population of surface-bound complexes. Molecular transduction converts and may amplify that state into a reporter, while physical observation determines which fraction becomes measurable information. The modules are replaceable, but the interface conditions—physical units, statistical background, time dependence, conservation and detection limits—are not optional.

The provisionally consolidated AR analysis provides the most developed mathematical demonstration of this architecture. It explains why identical equilibrium affinity can produce different kinetic and concentration responses, why buffer perturbations may alter apparent specificity without target-selective action, and why transport, finite inventory, time and receptor kinetics together govern microreactor performance. The TOI design-window and affinity-path analyses illustrate that extreme receptor affinity or extreme confinement need not be the most effective route to a better assay; such claims are specific to the stated operating conditions and remain subject to experimental validation.

In the full-chain ELISA and TOI examples, the same recognition inputs generate different AR populations, followed by distinct molecular reporter chemistries and observation mechanisms. A candidate conserved substrate-progress function now connects accumulating TMB product with instantaneous luminol emission while recognizing that delayed photon collection, direct fluorescence and multi-reactant chemistry require mechanism-specific interfaces. These cases illustrate the generality of the architecture without claiming that provisional T_m_ and T_s_ calibrations are final. Future refinement or substitution of those module models is expected. The decisive validation will be to determine each stage independently and predict the reported endpoint across new reagent amounts, reaction times and optical conditions without refitting the assembled chain. The enduring contribution sought by ART is thus a transferable mathematical language for how immunosensor components couple—not a particular expression chosen to represent one of them.

## Notes

### Competing Interest Statement

The authors have declared no competing interest.

### Funding

- Chinese Academy of Sciences

